# Phosphorylation of Gαi shapes canonical Gα(i)βγ/GPCR signaling

**DOI:** 10.1101/2022.09.11.507491

**Authors:** Suchismita Roy, Saptarshi Sinha, Ananta James Silas, Majid Ghassemian, Irina Kufareva, Pradipta Ghosh

## Abstract

A long-standing question in the field of signal transduction is to understand the interplay between distinct signaling pathways that control cell behavior. For growth factors and heterotrimeric G proteins, the two major signaling hubs in eukaryotes, the mechanisms of independent signal transduction have been extensively characterized; however, if/how they may cross talk remains obscure. Here we use linear-ion-trap mass spectrometry in combination with cell-based biophysical, biochemical, and phenotypic assays to chart at least three distinct ways in which growth factors may impact canonical Gα(i)βγ signaling downstream of a GPCR (CXCR4) via phosphorylation of Gαi. Phosphomimicking mutations in a cluster of residues in the αE helix (Y^154^/Y^155^) result in the suppression of agonist-induced Gα(i) activation while promoting constitutive Gβγ signaling; others in the P-loop (Ser^44^, Ser^47^, Thr^48^) suppress Gi activation entirely thus completely segregating the growth factor and GPCR pathways. While most phosphoevents appear to impact, as expected, the core properties of Gα(i) (conformational stability, nucleotide binding, Gβγ association and release, etc.), one phosphomimicking mutation promoted mislocalization of Gαi from the plasma membrane: a novel and unexpected mechanism of GPCR signal suppression. A phosphomutation of C-terminal Y^320^ was sufficient to orchestrate such suppression by protein compartmentalization. Findings not only elucidate how growth factor and chemokine signals crosstalk through phosphomodulation of Gαi, but also how such crosstalk may generate signal diversity.

## Introduction

A cell perceives and calibrates its responses to the external cues by a complex array of sensor proteins, commonly known as ‘receptors’; these receptors transduce signals from the plasma membrane (PM) to the cell’s interior via a cascade of downstream intermediates. Traditionally studied through reductionist approaches by dissecting a single cascade at a time, it is well-known that these distinct cascades crosstalk at multiple levels. Cross-talks between multiple cascades generate complex larger network for information flow, integration and diversification (through context, time and space), all of which ultimately coordinately controls cell fate (1–4).

In eukaryotes, two of the most widely studied signaling cascades are those that are initiated by the receptor tyrosine kinases (RTKs) and by the 7-transmembrane receptors coupled to heterotrimeric G proteins (GPCRs)(5). Upon sensing specific growth factors, RTKs auto-phosphorylate their cytoplasmic tails; subsequently, a variety of adaptor proteins dock onto those phosphosites to propagate the signal to the cell’s interior (6). Heterotrimeric (henceforth, trimeric) G proteins, on the other hand, serve as molecular switches, canonically acting downstream of GPCRs (7, 8). Agonist-bound GPCRs act as receptor guanine-nucleotide exchange factors (GEFs) for heterotrimeric G proteins, triggering GDP to GTP exchange on Gα subunit and releasing Gβγ dimers; GTP-bound Gα monomers and Gβγ dimers go on to bind the effector molecules and transduce a variety of signals (7).

Although it has been suggested that these two pathways crosstalk (9–13), in that G proteins may be activated downstream of RTK (14–24), whether and how these processes take place in cells, and their biological significance still remains ambiguous. Reports as early as the late 1980’s and early 1990’s had suggested that tyrosine phosphorylation of G proteins is one such mechanism (25–28); however, the identity of these sites and how they may impact G protein activity remained unknown until recently when Kalogriopoulos et al. (5) showed that multiple RTKs (but not non-RTKs) can directly phosphorylate and transactivate the α-subunit of the trimeric Gi protein. Upon growth factor stimulation, scaffolding of monomeric Gαi with RTKs (via a multi-modular signal transducer, GIV/a.k.a Girdin) (29) facilitates the phosphorylation on two tyrosines located within the inter-domain cleft of Gαi and one tyrosine located in the last β-strand (β6). Phosphorylation activates Gαi (by enhancing the basal exchange rate of GDP-to-GTP exchange) and enhances its ability to suppress cellular second messenger cAMP. These insights defined a tyrosine-based G protein signaling paradigm and revealed its importance in eukaryotes; they also raised at least two important questions—(i) Given the plethora of kinases that are activated upon growth factor stimulation (not just TKs), what other sites may be phosphorylated and how might they (alone or in combination) impact Gαi activation?; and (ii) How might these phosphoevents impact the ability of canonical GPCRs to transduce signals via Gi? Furthermore, although growth factors (EGF, epidermal growth factor and IGF1, insulin-like growth factor-1) were found to inhibit Gi coupling to and activation by GPCRs (CXCR4) (26, 30), mechanisms for such uncoupling/deactivation remain elusive.

Here we tackle these outstanding questions by interrogating how one prototypical growth factor/RTK system (EGF/EGF-receptor; EGFR) may shape one canonical Gi/GPCR system (the chemokine system, CXCL12/CXCR4) via phosphorylation of Gαi. We find that the phosphoevents inhibit different aspects of signaling and do so through a variety of mechanisms, with the location of the phosphosites clusters on the G protein being a key determinant of how such inhibition is orchestrated.

## Results

### Unique sites on Gαi are phosphorylated in the presence of epidermal growth factor (EGF)

In an earlier study (29), we demonstrated that Gαi3 undergoes tyrosine phosphorylation upon growth factor stimulation. Here, to determine which additional residues on Gαi are phosphorylated in the presence of growth factors, we conducted in-cell kinase assays, immunopurified the Gαi3-FLAG using an antibody against FLAG, and analyzed it by linear ion-trap mass spectrometry (MS) (31) (see *Methods* and **Figure 1a** for workflow). Prior to its use in MS studies, we confirmed the functionality and localization of the FLAG-tagged Gαi protein through a series of experiments addressing different aspects of Gαi function. In particular, we demonstrated that non-phosphorylated Gαi3-FLAG binds Gα-interacting protein (GAIP; **Supplementary Figure 1a**), a member of the RGS (regulators of G protein signaling) family, which serve as GAPs (GTPase-activating proteins) for Gαi (32); and that Gαi3-FLAG phosphotyrosination induced by EGF increases such binding. The C-terminal tag prohibits Gαi3-FLAG coupling to intact GPCRs; however, we showed that both non-phosphorylated and phosphotyrosinated Gαi3-FLAG bind the isolated third intercellular loop (icl3) of Lysophosphatidic Acid Receptor 1 (LPAR1) and alpha 2A adrenergic receptors (α2A, B) (**Supplementary Figure 1b**): icl3 is a region that ensures G protein specificity of receptor GEFs (33). The ability of non-phosphorylated and, to a lesser degree, phosphotyrosinated form to bind activator of G-protein signaling 3 (AGS3; **Supplementary Figure 1c**), a member of the GoLoco- (a.k.a. G-protein regulatory, GPR-) motif containing proteins that act as guanine nucleotide dissociation inhibitors (GDI) for Gαi (34, 35), was also demonstrated. We also confirmed the unaltered ability of Gαi3-FLAG to interact with Gβγ subunits in cells (**Supplementary Figure 1d**) and bind GIV, the prototypical member of the guanine nucleotide exchange modulators (GEM) (36, 37); **Supplementary Figure 1e**). Finally, the FLAG-tagged Gαi3 protein localized prominently at the plasma membrane in starved cells, with some cytosolic diffuse pattern observed upon EGF stimulation (**Supplementary Figure 1f-i**). The EGF/EGFR system was prioritized among other growth factors for these studies, not only because EGFR is the prototypical growth factor RTK, but also because it is the most widely studied RTK for pathway crosstalk with GPCRs (38–40).

**Figure 1.**
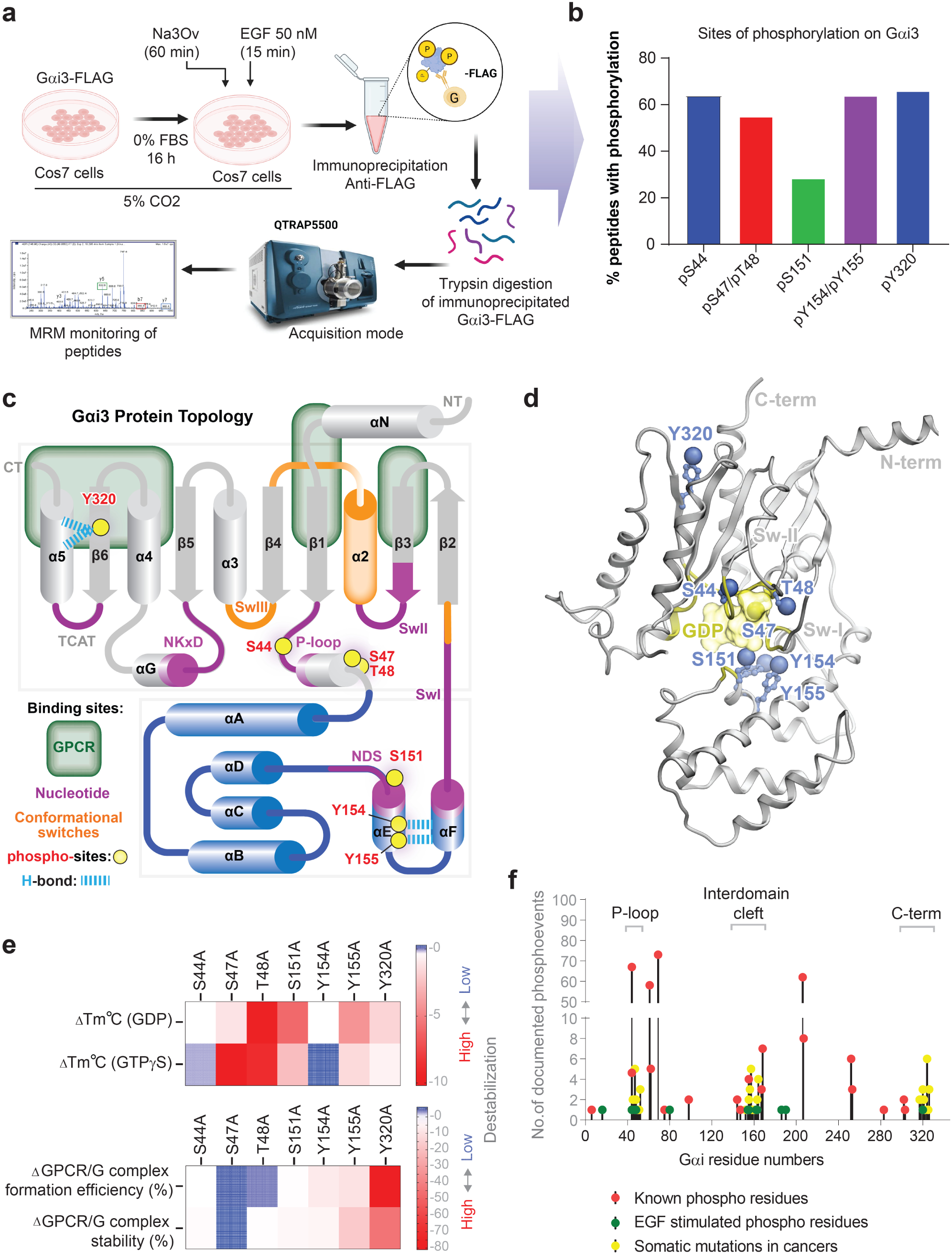
Analysis of Gαi3 phosphorylation in EGF-stimulated cells. a. Schematic of the experimental workflow used to discover phosphoevents on Gαi that occur in the presence of EGF. Briefly, serum-starved Cos7 cells co-expressing untagged EGFR and C-terminally FLAG-tagged Gαi3 were stimulated with 50 nM EGF. Gαi3 immunoprecipitated from cell lysates with FLAG mAb was digested on Protein-G beads and the tryptic fragments were analyzed using LC/MS without any further phosphoenrichment steps (see *Methods* for details). b. Bar graph displays the % phosphorylation of residues on Gαi3 (calculated as [phosphorylated peptides ÷ total detected peptides] x 100). c-d. Phosphosites on Gαi3 that were identified by LC/MS in a-b are projected on a topology map of the Gαi3 protein (modified from (76) with conformational switches and binding sites of key interactors marked (**c**) and on a ribbon diagram of the solved crystal structure of Gαi (PDB: 5tdh; **d**). Residues indicated in red highlight phosphosites which were mutated to phosphomimic (or non-phoshorylatable) aa in this work. e. Heatmaps display the experimentally determined changes in the thermal stability of Gαi1 and its ability to form GPCR complexes upon mutating key phosphosites (identified in a-b) on into Ala(A) reported by a prior work (41). The thermal stability of each mutant is displayed (from top to bottom) in the inactive GDP-bound, the active GTP(GTP*γ*S)-bound states, and the efficiency of formation (relative abundance) and relative stability of the reconstituted Rhodopsin-G_i_ protein complex. Blue and red colors indicate low and high degrees of destabilization. Source data is presented in **Supplemental Data File 3**. f. Lollipop diagram shows all documented S/T/Y phosphorylation events on Gαi1, Gαi2, and Gαi3 and somatic mutations in cancers (red: data from Phosphosite.org; Green: Our experimental data; Yellow: Data from COSMIC; https://cancer.sanger.ac.uk/cosmic).

Two independent MS analyses were performed; each time, samples were processed without any phosphoenrichment to allow for the assessment of the relative stoichiometry of the various phosphosites. Different sets of phosphorylated peptides were identified (**Figure 1b**); 7 sites were prioritized based on their confidence score, location within the G protein and corroboration in independent LC/MS studies (see **Table 1**).

**Table 1.**
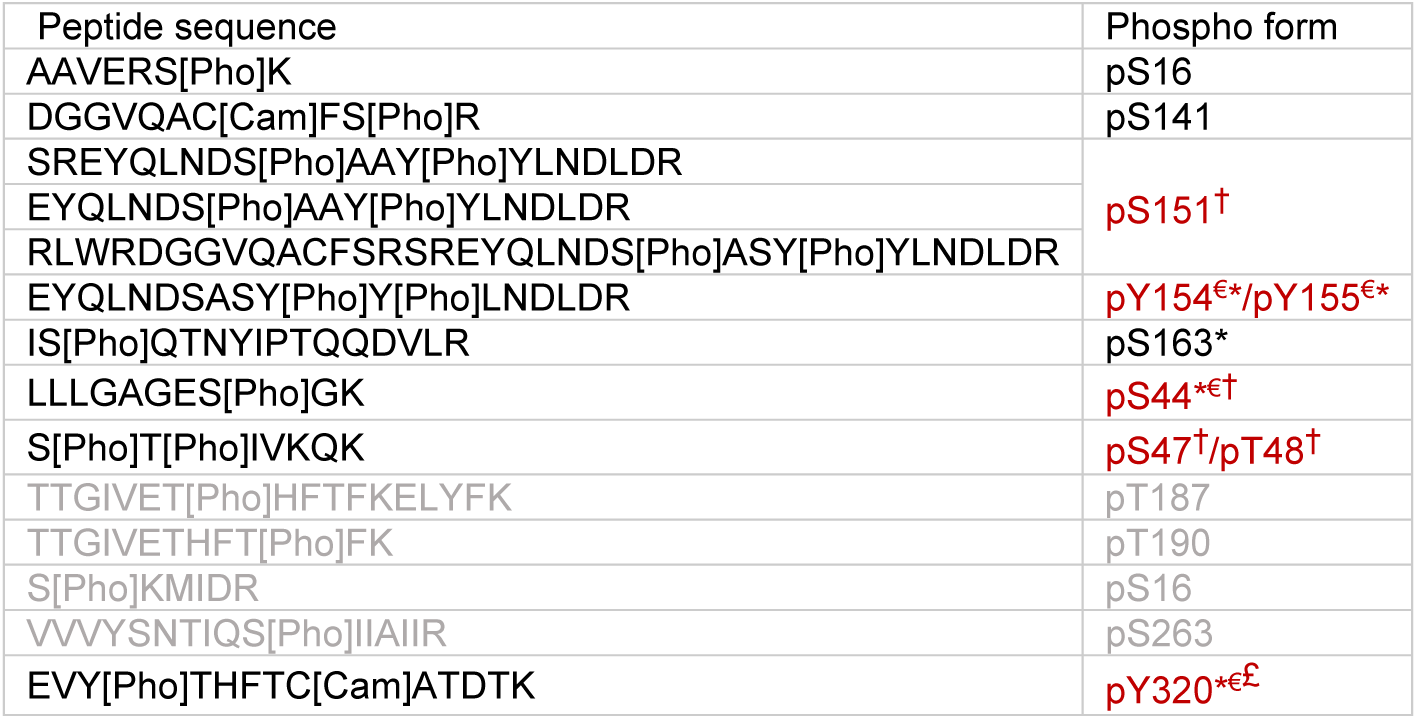
Phosphosites identified by linear ion-trap mass spectrometry. Red font indicates the sites studied in this work. * Sites reported as phosphorylated in phosphosite.org; € Sites reported previously to be phosphorylated in ligand-dependent manner; † Sites neighboring nucleotide-binding pocket; £ Sites facing GPCR-binding interface.

Projection of these residues on a topology map of Gαi3 (**Figure 1c**) and a solved crystal structure of the highly homologous Gαi1 (**Figure 1d**) revealed that phosphorylated positions are located in some of the key regions within Gαi, spanning both all-helical (AHD) and Ras-like domains, the nucleotide binding site, and the interdomain cleft. For example, Ser^44^, Ser^47^ and Thr^48^ are in the phosphate-coordinating P loop, Ser^151^ and Y^154^/Y^155^ are in the αE helix where they stabilize the αF helix and the conformational Switch-I (Sw-I), and Y^320^ is in the β6 strand where it interacts with the C-terminal α5 helix directly engaging with activated GPCRs (41). Ser^44^, Ser^47^, Thr^48^ and Y^154^ are strictly conserved across all mammalian Gα proteins, whereas S^144^ is replaced by Arg, Ala, and Asn, S^151^ by Asn and Cys, and both Y^155^ and Y^320^ by Phe in selected Gα protein subtypes (**Supplementary Figure 2**). Barring two recently reported tyrosine (Y^154^/Y^155^) that are phosphorylated by RTKs within the interdomain cleft of Gαi (5), the other sites remain uncharacterized and their importance unknown.

Previously, the role of Gαi residues in the formation and stability of nucleotide-bound Gαi states or GPCR(rhodopsin)•Gαi complexes has been characterized comprehensively at a single amino-acid (aa) level by alanine (Ala; A) scanning mutagenesis (41). A limited re-analysis of the published dataset from (41), focused exclusively on the residues of our interest, revealed that both P-loop residues Ser^47^ and Thr^48^ are critical for GTP binding, with Thr^48^ also being important for GDP binding, whereas mutations in Ser^44^ do not have much impact on either state (**Figure 1e***-top*). Ala mutations of the interdomain cleft residues Ser^151^ and Y^155^ destabilized the GDP-bound state to a greater extent than the GTP-bound state; while elimination of Y154 did not have much impact on either state (**Figure 1e***-top*). The C-terminally located Y^320^ was required for GPCR•Gαi complex formation and stability (**Figure 1e***-bottom*); however, Ala substitution at this site had only a moderate effect on the nucleotide-bound states (**Figure 1e***-top*) and no effect on heterotrimer formation (41). These findings suggest that phosphorylation at residues located in the P-loop and those buried within the interdomain cleft may primarily alter the kinetics of nucleotide exchange on the Gαi protein (and consequently its activity, conformation, and ability to sequester or release Gβγ), whereas phosphorylation at the C-terminal Y^320^ may primarily impair receptor engagement/recruitment.

Finally, the unique phosphosites identified in the presence of EGF were compared to phosphoevents on Gαi1-3 previously detected by high throughput mass spectrometry (HTP-MS) (21, 23, 24, 26, 42, 43) [cataloged at phosphosite.org], and with previously reported sites of somatic mutations in cancer [cataloged in the COSMIC database]. Such analysis showed that, for 4/7 phosphosites that we identified, the previously reported phosphoevents were induced by ligands (see **Table 1**), and that other previously observed phosphoevents and mutations tend to cluster in the same three regions of the Gαi protein: P-loop, interdomain cleft, and the C-terminus, implying that these could represent ‘hot spots’ within the allosteric switch (**Figure 1f**).

Taken together, the phosphoevents we selected to study further in this work (**Figure 1b**; **Table 1**), are either reported to be ligand-dependent (Ser^44^, Y^154^, Y^155^, Y^320^) or are likely to be so, given their role in coordinating nucleotide binding in the dephosphorylated state (Ser^47^, Thr^48^, Ser^151^). We hypothesized that phosphomodifications that we observe in growth factor stimulated cells may have interesting and meaningful consequences for canonical heterotrimeric G protein signaling.

### Study design and rationale

To study the impact of phosphorylation on G protein signaling, we resorted to the well-accepted norm of comparing the WT protein (that is expected to reversibly undergo cycles of phosphorylation and dephosphorylation as expected in physiologic state) against mutants with phosphomimicking and non-phosphorylatable substitutions that ‘lock’ the function of the protein in constitutively phosphorylated and dephosphorylated states, respectively. We generated phosphomimicking (S→D; T→E, Y→E), and in the case of Y^320^, non-phosphorylatable (Y→F) mutants of these residues (see **Table 2**) based on the standard algorithm for residue-specific phosphorylation site replacements (44). Because Ala substitutions at the identified Ser/Thr (but not the identified Tyr) sites strongly impact the stability and properties of Gαi (**Figure 1e)** (41), the use of Ser/Thr → Ala substitutions were deemed inappropriate; the inherent effect of such substitution on protein stability was expected to confound any other effect of constitutive dephosphorylation in ways that would be impossible to untwine. Our decision to make them as single mutations, or as combinations, was guided by their location, concomitantly phosphorylated state in cells (**Figure 1b**), and by theme, that is, pan-tyrosine, Y^154^, Y^155^ and Y^320^ combined or pan-pSer/Thr/Tyr, in which all sites identified and prioritized in this work were simultaneously substituted within the same construct (see list of mutants in **Table 2**). To enable functional characterization without the interference of endogenous G protein, all mutations were generated in a Gαi3 construct in which a cysteine (Cys; C) at position 351 was mutated to isoleucine (Ile; I): this C351I mutation renders the G protein insensitive to pertussis toxin (PTX) (45, 46) and allows, in conjunction with PTX treatment, to selectively assess the GPCR-related functions of the mutant G protein (47). Consequently, in all our assays, the PTX-sensitive wild-type Gαi3 (WTs) and the PTX-resistant wild-type Gαi3 (WTr) served as negative and positive controls, respectively.

**Table 2.** Inventory of constructs and the annotation used in this work.

| Constructs | Annotation | Comments |
| --- | --- | --- |
| *Gai3* | WTs | PTX sensitive |
| *Gai3 (C351I)* | WTr | PTX resistant (working as backbone control for all the mutants) |
| *Gai3-S44D(C351I)* | S44D |  |
| *Gai3-S151D(C351I)* | S151D |  |
| *Gai3-Y320E(C351I)* | Y320E |  |
| *Gai3-Y320F(C351I)* | Y320F |  |
| *Gai3-S47D-T48E(C351I)* | S47D-T48E |  |
| *Gai3-YY154-155EE(C351I)* | YY154-155EE |  |
| *Gai3-YYY154-155-320-EEE(C351I)* | YYY154-155-320EEE |  |
| *Gai3-YYY154-155-320-EEF(C351I)* | YYY154-155-320EEF |  |
| *Gai3-S44D-S47D-T48E-Y154E-Y155E(C351I)* | Pan-pS/T/Y |  |

To assess the impact of the phosphomimicking Gαi mutations on canonical signaling of Gi coupled seven-transmembrane receptors (further abbreviated as *GiPCRs* for “*Gi protein-coupled receptors*”), we used the prototypical family member, chemokine receptor 4 (CXCR4), which couples to and transmits its signals primarily through the PTX-sensitive G_i_ proteins (48). We chose CXCR4 because of the well-studied Gi-mediated signaling pathways that it activates in response to its endogenous chemokine agonist CXCL12, and because of its importance in cancer invasion and metastasis (48); the latter is a process that is enriched in pathway crosstalk between growth factors and GPCRs (38). In the case of CXCR4, prior studies have alluded to the presence of pathway crosstalk with the EGF/EGFR pathway and role of such crosstalk in promoting tumor progression (49, 50).

As a model system for our studies of CXCR4-driven Gαi3 signaling, we chose HeLa cells because of abundant endogenous CXCR4 (51) and their responsiveness to CXCL12 (51, 52). HeLa are also known to express the second receptor for CXCL12, ACKR3 (53–55); however, as an *atypical* chemokine receptor, ACKR3 exclusively transmits its signals through β-arrestin (53, 56–58) and therefore is unlikely to confound our assessment of Gαi mutant functionality.

### Impact of the Gαi phosphomimicking mutations on canonical GiPCR-mediated Gβγ signaling

First we assessed Gβγ release from the receptor-activated activated heterotrimeric Gαi/Gβγ complex, which is one of the earliest steps and a key event in the transduction of GiPCR signals (59, 60). We assessed Gβγ release and activation using agonist-induced changes in BRET (bioluminescence resonance energy transfer) between mVenus-tagged Gβγ (BRET acceptor) and RLuc8-tagged C-terminus of G protein-coupled receptor kinase- 3 (GRK3ct) (BRET donor) (61–63) (**Figure 2a-b**). A CXCL12-induced increase in BRET ratio indicates dissociation of the Gαi/Gβγ heterotrimer and release of free Gβγ, which then becomes available for interaction with GRK3ct. Moreover, prior to CXCL12 stimulation, a higher basal BRET ratio between Gβγ and GRK3ct indicates basal excess of free (non-Gαi-bound) Gβγ dimers, likely due to the compromised ability of mutant Gαi to sequester them and to form stable Gαi/Gβγ heterotrimeric complexes.

**Figure 2.**
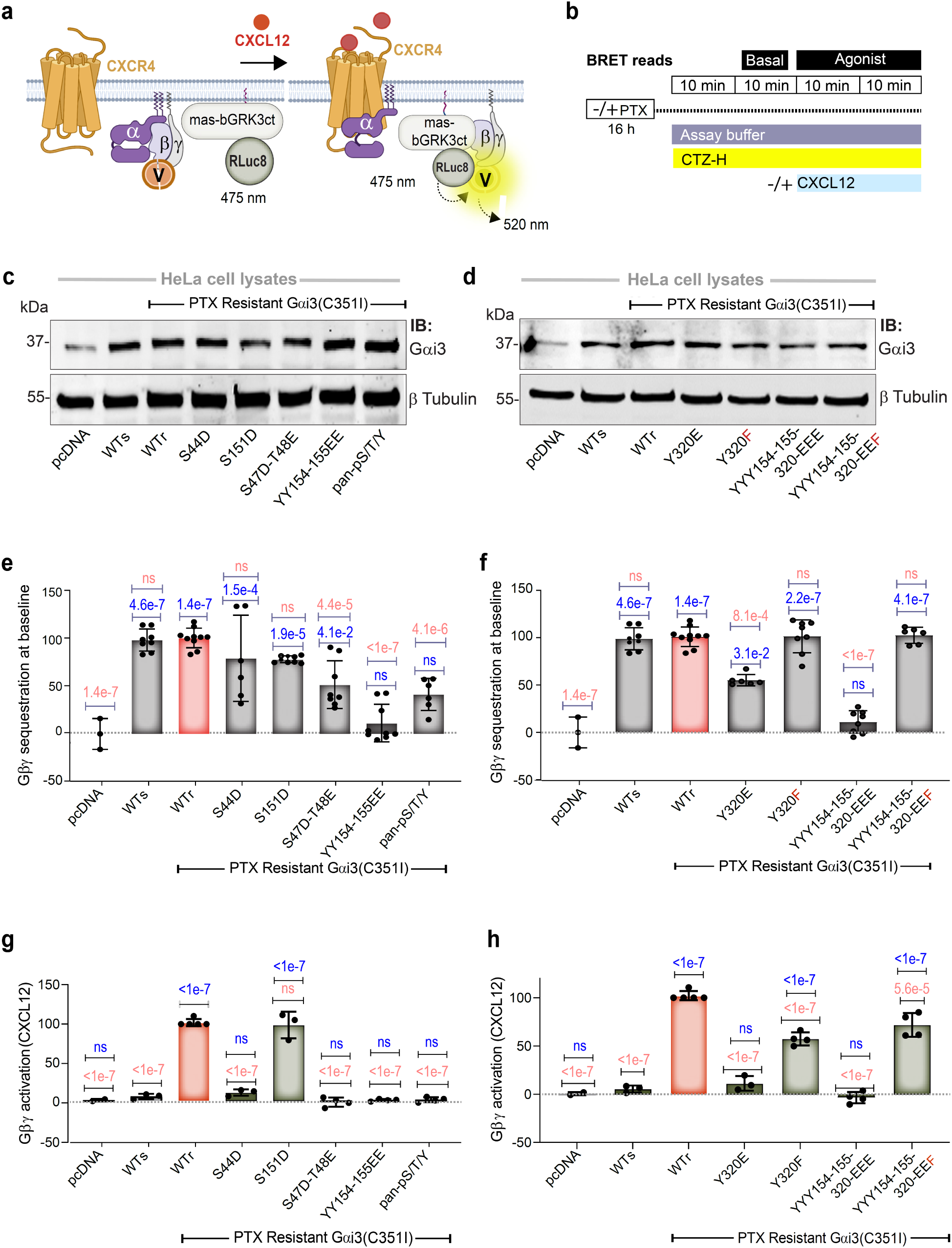
Effect of phosphomimicking mutations in Gαi3 on basal and CXCL12- stimulated Gβγ activation downstream of CXCR4. a-b. Schematics of the principles (a) and the timeline for various treatments and readouts (b) of the BRET-based assay for assessing basal and ligand-stimulated Gβγ release from trimeric Gi. Prior to ligand stimulation (basal), Venus-tagged Gβ1/Gγ2 complexes bind Gαi●GDP to form inactive trimers. Upon addition of CXCL12 (100 nM), nucleotide exchange triggers the release of Venus-Gβγ dimers from Gαi which allows them to bind GRK3ct- RLuc8 and results in an increase of the BRET signal. c-d. Immunoblots showing the abundance of each phosphomimicking (or non-phosphorylatable) Gαi mutant in equal aliquots of HeLa cell lysates (∼35 μg of total protein, as estimated by Bradford method). β-Tubulin was used as a loading control. e-f. Bar graphs display the % Gβγ sequestration observed in the case of each mutant at baseline, before CXCL12 stimulation, all analyzed under the same conditions as outlined in b, but in two sets as represented in the graphs. Results are expressed as the **suppression** of basal mVenus-Gβγ/GRK3ct-RLuc8 association normalized to a set scale of 0-100%, where 0% is in the absence of transfected Gαi (pcDNA, negative control, no suppression) and 100% is in the presence of either PTX-sensitive (WTs) or PTX-resistant (WTr) Gαi3 (positive controls, both maximally suppressing the basal mVenus-Gβγ/GRK3ct-RLuc8 BRET). All assays were performed in the PTX- treated condition. g-h. Bar graphs display the % Gβγ activation upon CXCL12 stimulation observed in the case of each mutant in **Table 2**, all analyzed under the same conditions as outlined in b, but in two sets as represented in the graphs. Results are expressed as ligand-induced increase in mVenus-Gβγ/GRK3ct-RLuc8 association (indicative of Gβγ release from Gαi and resulting activation), and normalized observed to a set scale of 0-100%, where 0% is pcDNA and the PTX-sensitive Gαi3 (WTs) (negative controls, unable to release and activate Gβγ in response to CXCL12 in PTX-treated conditions) and 100% is the PTX-resistant Gαi3 (WTr, positive control, maximum release and activation of Gβγ in response to CXCL12 in PTX-treated condition). *Statistics*: *P* values were determined by one-way ANOVA with Tukey’s multiple comparison’s test. P Values in blue and red fonts indicate significance compared to negative control (pcDNA/WTs) and positive controls (WTr), respectively. Error bars represent ± S.E.M; n=4 independent experiments, each with 3 technical replicates. In panels (e-f), PTX-treated and non-PTX treated samples from the same day were used as separate biological replicates thus producing n=8, each with 3 technical replicates. See also **Supplementary** Figure 3 for the composite data for all conditions.

After confirming that exogenous transfection of various Gαi mutant constructs in HeLa cells results in comparable expression (**Figure 2c-d**), BRET was assessed with or without PTX pretreatment and with or without CXCL12 stimulation. Prior to CXCL12 stimulation, high Gβγ/GRK3ct BRET was observed in the absence of exogenously transfected Gαi (labeled “pcDNA”, 0% suppression of basal Gβγ/GRK3ct association), this BRET was strongly suppressed when either WTs or WTr Gαi was co-transfected (100% suppression of basal Gβγ/GRK3ct association, **Figure 2e-f**), indicating the ability of both forms to sequester Gβγ and form stable heterotrimeric complexes, unaffected by PTX treatment. In the assessment of CXCL12-induced increases in Gβγ/GRK3ct association, as expected, WTs was sensitive to PTX treatment, whereas WTr remained unaffected (compare CXCL2 vs CXCL12 + PTX; **Supplementary Figure 3a-b**), indicating that the PTX-treated conditions are appropriate for drawing further conclusions regarding the impact of our mutant constructs without interference from endogenous Gαi.

The three variants containing phosphomimicking modifications of Y^154^ and Y^155^ (YY154-155EE, YY154- 155EE/Y320E, and pan-S/T/Y) demonstrated significant decreases in their ability to suppress the basal Gβγ/GRK3ct BRET signal when compared to the positive control WTr, and no differences when compared to the negative control (pcDNA, no transfected Gαi) (**Figure 2e-f**; **Table 3***-column #3*), indicating that these mutants are severely impaired in Gβγ binding. These mutants also failed to show an increase in Gβγ/GRK3ct association (indicative of Gβγ release from Gαi) upon ligand stimulation (**Figure 2g-h**; **Table 3***-column #4*), consistent with their inability to bind Gβγ in the first place. By contrast, the P-loop mutants were either proficient (S44D) or partially proficient (S47D-T48E) in Gβγ sequestration at baseline (p values of 1.5e-4 and 4.1e-2 respectively, when compared to the negative control pcDNA; and p values of n.s and 4.4e-5, respectively, when compared to the positive control WTr); however, they were still defective in Gβγ release upon ligand stimulation (both not significantly different from negative control WTs; p value <1e-7 for both S44D and S47-T48E when compared to the positive control WTr). This may be a consequence of the inability of these mutants to bind GTP: agonist-induced Gβγ release requires that Gαi is GTP-bound, but for at least two other substitutions at position 47 of Gαi (Ala and Asn), severe impairment of GTP binding has been previously reported (41, 64), with S47N also known to preferentially stabilize (or trap) a nucleotide-free, GPCR-bound state of the Gαi/Gβγ heterotrimer (64–66). Additionally, an Ala substitution at position 48 has been shown to abrogate the binding of both GDP and GTP (59). As for the C-terminal phosphosite Y320, the phosphomimic Y320E substitution was partially deficient in Gβγ sequestration and incapable of Gβγ release after ligand stimulation. By contrast, the Y320F mutant was fully and partially proficient in Gβγ sequestration and release, respectively, suggesting that phosphorylation of Gαi at Y^320^ may be sufficient to inhibit canonical Gi signaling. Interestingly, although the double YY154-155EE mutant and the triple YYY154-155-320EEE mutant were both inert in basal Gβγ sequestration (**Figure 2f**) and ligand stimulated Gβγ release (**Figure 2h**), a replacement of Y^320^ by the non-phosphorylatable “F” led to a mutant (YYY154-155-320EEF) that regained full proficiency in both Gβγ sequestration (p value = 4e-7 compared to negative control, and not significantly different from positive control) and partial proficiency in ligand-stimulated Gβγ release (p value <1e-7 and 5.6e-5 compared to negative and positive controls, respectively). This indicates that the Y320F substitution rescued the defects observed in the double Y→E mutant within the interdomain cleft (*p* value <1e-7 for both basal sequestration and ligand-induced release; see **Table S1-S2; Table 3***-column #3- 4*).

**Table 3.**
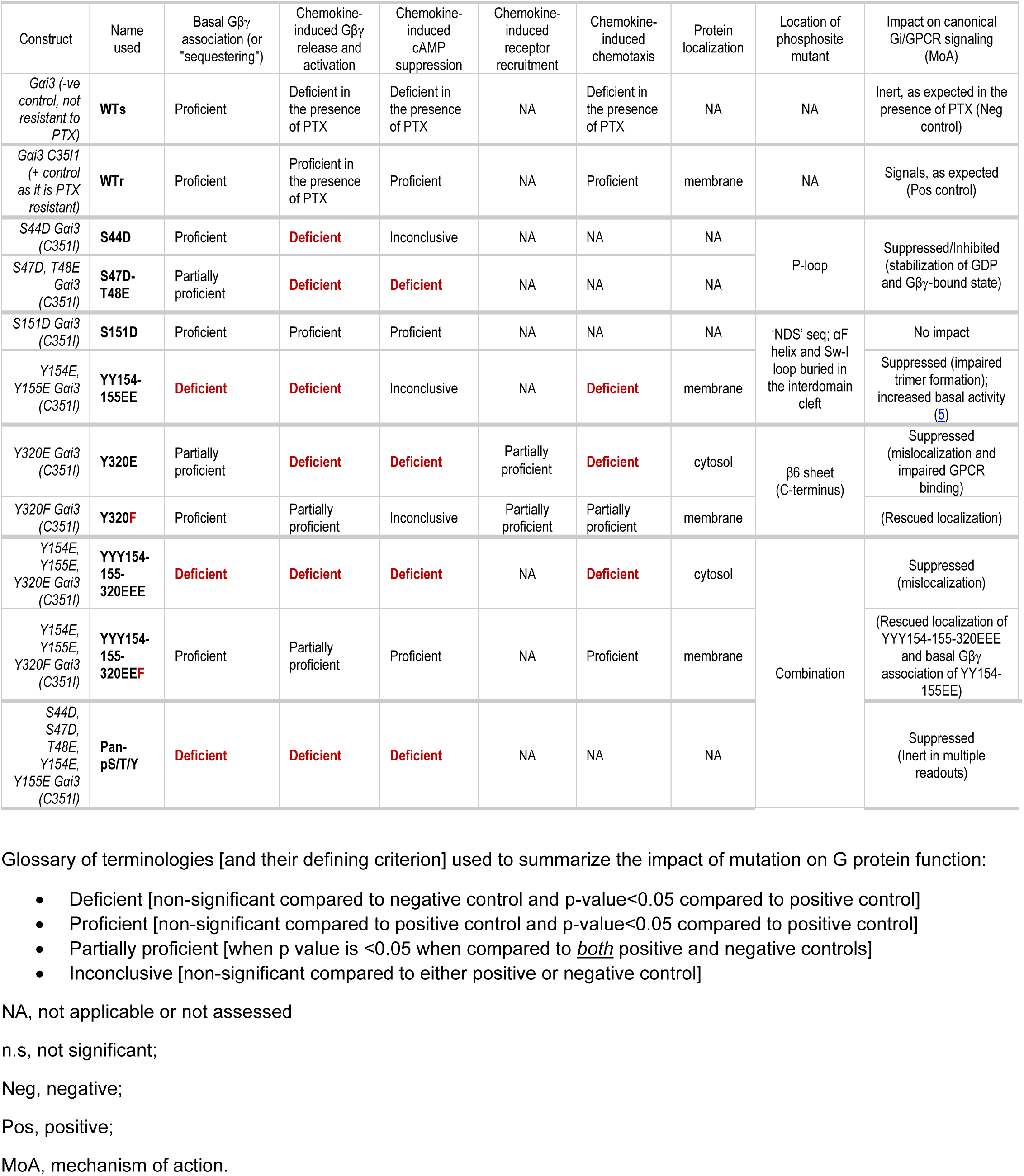
Summary of experimentally derived findings.

These findings suggest that Gαi phosphomodifications that we observed in growth factor-stimulated cells can modulate the canonical Gβγ signaling in at least two ways: (i) impair the ability of the Gαi to form trimers with Gβγ at basal state, and consequently, inhibit canonical ligand-induced signaling while allowing for constitutive activation of Gβγ (exemplified by phosphomimicking mutations at the interdomain cleft and at the C-terminus); (ii) inhibit ligand-stimulated Gβγ release but preserve, partially or fully, the ability to sequester Gβγ and form trimers at basal state (such as Ser/Thr phosphorylation in the P-loop). The mutation mimicking pan-phosphorylation at all Ser/Thr/ Tyr sites follows the first pattern. Intriguingly, phosphorylation at Y^320^ (mimicked by E) was sufficient to inhibit signaling, regardless of whether it was assessed independently by comparing Y320E vs Y320F (*p* value 0.0014 for Gβγ sequestration and 7.0e-7 for Gβγ release, respectively; **Table S1-S2**) or in combination with other phosphotyrosines by comparing YYY154-155-320EEE vs YYY154-155-320EEF (*p* values <1 e-7 for both basal Gβγ sequestration and ligand-induced Gβγ release; **Table S1-S2; Table 3***-column #3-4*). Even more interesting, the non-phosphorylatable Y320F appears to rescue defects caused by the double YY154-155EE mutation.

### Impact of the Gαi phosphomimicking mutations on canonical GiPCR-triggered cyclic AMP inhibition

As a G_i_ protein-coupled chemokine receptor (GiPCR), CXCR4 inhibits adenylyl cyclase activity and decreases intracellular cAMP (67); we asked what, if any, impact the Gαi phosphomodifications have on this key step in chemokine signaling. To this end, we used the established BRET-based sensor for cAMP, CAMYEL (cAMP sensor using YFP-Epac-RLuc), which can monitor intracellular concentrations of cAMP in real time in live cells (68) (**Figure 3a-b**). By the design of the sensor, a decrease in BRET ratio (acceptor emission divided by donor emission) corresponds to an increase in cAMP; therefore, for an intuitive display, in **Supplementary Figure 4b**, we expressed results of this assay as inverse BRET (donor emission divided by acceptor emission); for **Supplementary Figure 4a** and **Figure 3c-d**, this quantity was further converted into cAMP suppression. Cyclic AMP levels were assessed following the addition of forskolin (FSK) to cells pre-treated (or not) with PTX and stimulated (or not) with CXCL12 (**Figure 3b**; *see Methods*). Basal suppression or CXCL12-induced suppression of FSK-induced cAMP were computed as % compared to negative control [the PTX-sensitive wild-type Gαi3 (WTs)] set at 0% and the positive control [PTX-resistant wild-type Gαi3 (WTr)] set at 100%. Compared to the positive control WTr, none of the mutants significantly impacted the basal suppression of FSK-induced cAMP (**Supplementary Figure 4a**). In chemokine-induced cAMP suppression, the P-loop (Ser^47^/Thr^48^), the β6-strand (Y320E), the pan-Y (YYY154-Y155EEE), and the pan-S/T/Y mutants were significantly different from the positive control and not different from the negative control (**Figure 3c-d**; **Supplementary Figure 4b**; **Table 3***-column #5*), indicating that these mutations abrogated CXCL12-induced cAMP suppression by Gαi. This was similar to their impairment in ligand-induced Gβγ release. By contrast, S151D and YYY154-155-320EEF were no different from the positive control and significantly different from the negative control, indicating that they retained the cAMP suppression ability. The rest of the mutants (S44D, YY154-Y55EE, and Y320F were not significantly different from either positive or negative control so that no conclusions could be made about their impact on the cAMP suppression.

**Figure 3.**
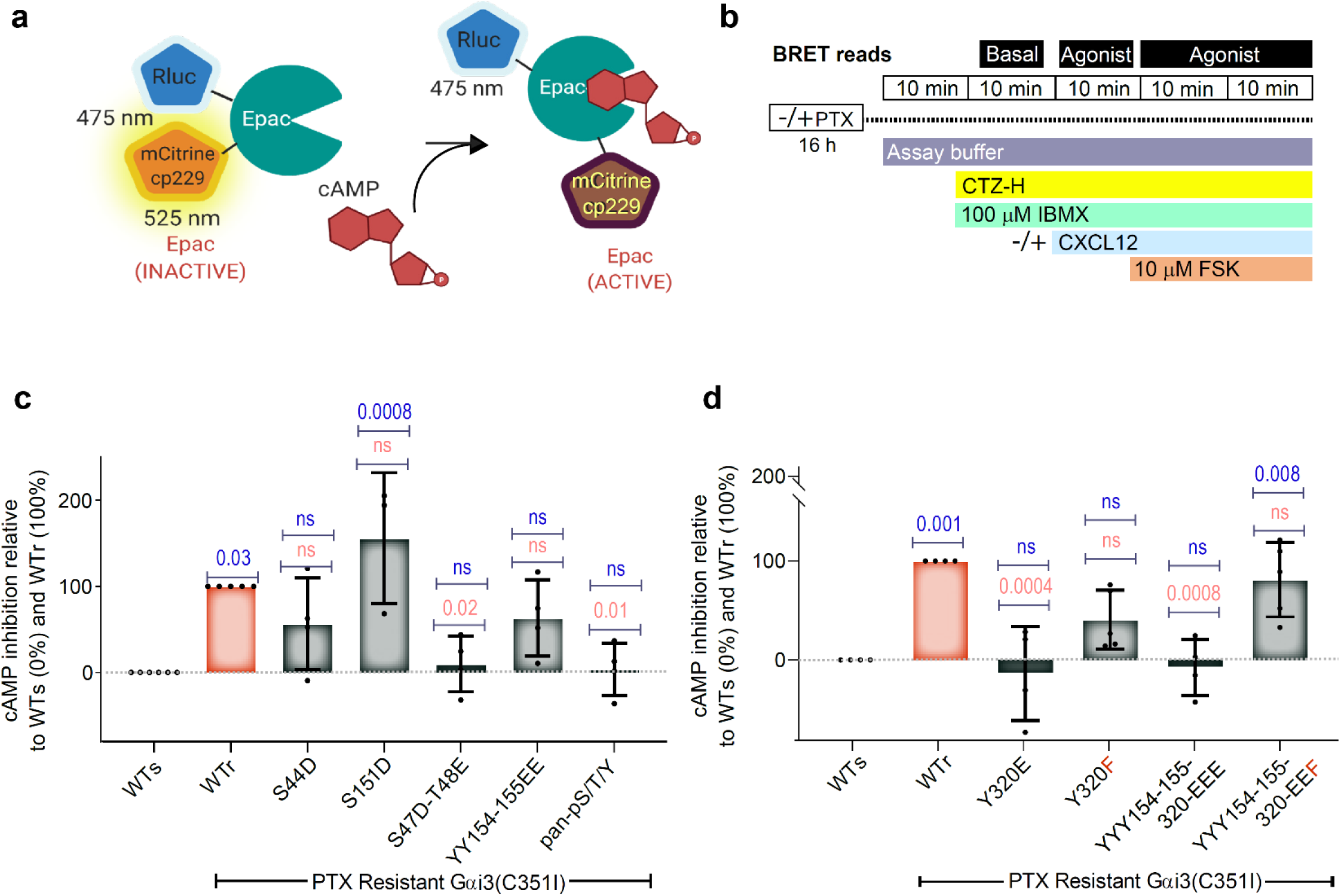
Effect of phosphomimicking mutations in Gαi3 on CXCL12-stimulated cAMP suppression downstream of CXCR4. a-b. Schematics of the principle (a) and the timeline for various treatments and readouts (b) for the BRET-based CAMYEL (*<u>cAM</u>*P sensor using *<u>Y</u>*FP-*<u>E</u>*pac-R*<u>L</u>*uc) assay for assessing cAMP suppression by WT and mutant Gαi. Forskolin (FSK) is used to increase cellular cAMP, which, upon binding to CAMYEL, decreases the BRET signal. Activation of Gαi suppresses the extent of FSK-stimulated cAMP production, and consequently, increases the BRET signal. c, d. Bar graphs display the % CXCL12-triggered (100 nM) inhibition of FSK-induced cAMP in HeLa cells transiently expressing the indicated Gαi variants and pre-treated with PTX. Results are normalized to a set scale of 0-100%, where 0% is cAMP inhibition mediated by the PTX-sensitive Gαi3 in PTX-treated conditions (WTs, negative control, no cAMP inhibition) and 100% is cAMP inhibition mediated by the PTX-resistant Gαi3 (WTr, positive control, maximum inhibition). *Statistics*: *P* values were determined by one-way ANOVA with Tukey’s multiple comparison’s test. P-Values in blue and red fonts indicate significance compared to the negative control wild-type sensitive (WTs) and positive control resistant (WTr) G protein, respectively. Error bars represent ± S.E.M; n=4 independent experiments, each with 3 technical replicates. See **Supplementary** Figure 4a for the impact of each Gαi3 mutant on the basal FSK-stimulated cellular cAMP levels and **Supplementary** Figure 4b for each biological replicate.

These findings suggest that some of the Gαi phosphomodifications that we observed in growth factor-treated cells can impair the ability of the Gαi to suppress cAMP downstream of agonist-activated GiPCRs, as exemplified by phosphorylation at Ser^47^ and Thr^48^ in the P-loop or the Y^320^ phosphorylation at the C-terminus. Other modifications are predicted to exert no effect, as exemplified by a selected phosphoserine-mimicking mutation in the interdomain cleft, Ser^151^. Intriguingly, the phosphomimicking mutation at Y^320^ (Y320E) is sufficient to inhibit signaling, whereas the combination of the non-phosphorylatable Y320F mutant with YY154-155EE does not appear to be significantly compromised (**Table 3***-column #5*).

### Phosphomimicking and non-phosphorylatable mutations at Y^320^ impact Gαi recruitment to the receptor

The observed profound and distinct effects of the phosphomimicking mutation Y320E and non-phosphorylatable mutation Y320F on the canonical signaling of Gαi are consistent with the location of Y^320^ in the Gαi β6 strand, where it directly contacts the GiPCR and the receptor-engaging C-terminal α5 helix of Gαi (41, 69) (**Figure 4a- b**), the latter through both non-polar interactions and hydrogen bonds. Because if this, we asked if/how the Y320E/F mutants impact coupling of Gαi to GPCR. To assess receptor recruitment, we used a well-defined BRET-based assay in which Venus-tagged engineered “mini” Gα proteins engage RLuc-tagged receptors (70).

**Figure 4:**
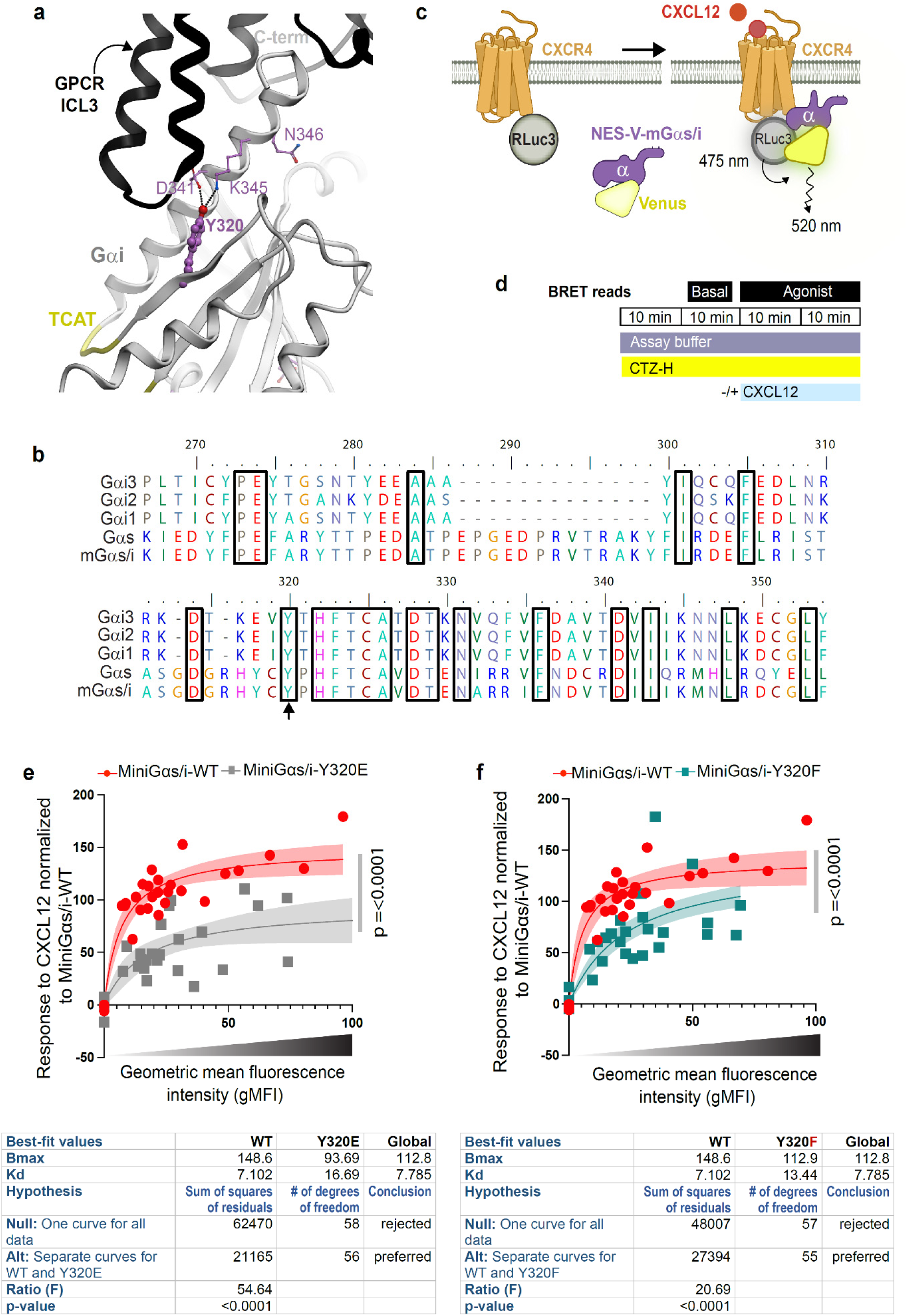
Impact of phosphomimicking and non-phosphorylatable mutants of Y^320^ on GiPCR recruitment to Gαi. a. A ribbon diagram of the solved crystal structure of Gαi-bound GPCR (PDB: 6cmo) highlights the interaction surface of the GiPCR intracellular loop 3 (ICL3) with the C terminal α5 helix of Gαi. Residues crucial for the stabilization of the receptor-bound state are interactions between the β6 (Y320) and the α5 (D341, K345 and N346). b. An alignment of rat Gαi1-3 sequences with Gαs and the mini-Gαs/i chimera used here is shown. Arrow highlights the residue Y320. c-d. Schematics of the principle (c) and the timeline for various treatments and readouts (d) of the BRET-based assay for assessing GPCR recruitment to the WT and mutant mini- Gαs/i chimeras. Proximity between the receptor (tagged with the BRET donor, Renilla luciferase 3) and the mini-Gαs/i (tagged with the BRET acceptor, Venus) results in energy transfer (see *Methods*). e-f. Shown are the results of BRET based saturation assays for mini-Gαs/i-WT vs mini-Gαs/i-Y320E (e) and mini-Gαs/i-WT vs mini-Gαs/i-Y320F (f) when a fixed amount of donor DNA (100 ng CXCR4-RLuc3 per well in a 6-well plate) is co-transfected with increasing amounts of the acceptor DNA (mVenus mini-Gαs/i chimera; ranging from 0 ng to 2500 ng of DNA per well in a 6-well plate). The x axis indicates geometric mean fluorescence intensity (gMFI), whereas the y axis displays CXCL12 response normalized to mini-Gαs/i-WT (positive control). N = 5 independent experiments, each with 4 technical replicates. The semi-transparent bands associated with each fitting curve represent the range of 95% confidence intervals (CI). F test has been performed to compute the significance of separate fits vs global fit (see **Table S4**). Bar graphs showing the CXCL12 response (normalized to mini-Gαs/i-WT), for the increasing amount of the acceptors, are presented in **Supplementary** Figure 5a. Lineweaver-Burk (LB) plots (**Supplementary** Figure 5b**, c)** and the accompanying statistical tests (**Table S5**) are provided. A representative flow cytometry quantification for acceptor expression is shown in **Supplementary** Figure 5d-f. *Statistics*: To calculate the Kd and Bmax we have used one site binding hyperbolic model fitting. Pairwise comparison between data specific fit and global fit within mini-Gαs/i-Y320E and mini-Gαs/i-Y320F, using extra sum-of-squares F test has been performed on both the model parameters (see **Table S4**).

Mini Gα proteins lack membrane anchors, N-terminal Gβγ-binding surface, and the α-helical domain (AHD), and contain a nuclear export signal (NES) (70). Ligand-induced coupling of a miniGα to an Rluc3-tagged GPCR results in higher BRET ratio (**Figure 4c, d**). For Gi coupled receptors, the appropriate miniGα is miniGαs/i, a chimera between Gαs and Gαi. We confirmed by sequence alignment that Y^320^ in Gαi is conserved in Gαs, and that the engineered miniGαs/i protein also contains a Y(Tyr) at the corresponding position (**Figure 4b**); the corresponding Y(Tyr) in miniGαs/i was then mutated to either the phosphomimicking E(Glu) or the non-phosphorylatable F(Phe) amino acid. We used these mutants alongside the WT miniGαs/i in a BRET-based acceptor saturation experiment, in which the transfected amount of the donor (CXCR4 C-terminally tagged with RLuc3) is kept constant while the amount of the acceptor (Venus-tagged miniGαs/i-WT or the Y320E/F mutants) is titrated. The resulting normalized BRET ratio showed that, as expected, GPCR coupling upon ligand stimulation gets saturated in the case of miniGαs/i-WT, but is impaired to different degrees in the case of the Y320E/F mutants (**Figure 4e-f**). While the Y320E mutant is severely impaired (according to F-test, null hypothesis a.k.a. global fit is rejected with p-value < 0.0001; mutant Kd=16.69 vs WT Kd=7.1, mutant Bmax=93.69 vs WT Bmax=148.6; **Figure 4e, Table S4**), the Y320F mutant’s functionality is intermediate (null hypothesis also rejected with p-value < 0.0001, mutant Kd=13.44, mutant Bmax=112.9; **Figure 4f, Table S4**). This finding was consistent with observations across all tested acceptor transfection levels (**Supplementary Figure 5a**) and was further confirmed through double-reciprocal (Lineweaver-Burk, LB) plot analysis (**Supplementary Figure 5b, c, Table S5**): the slopes of the LB plots were significantly non-zero for mini-Gαs/i- WT and mini-Gαs/i-Y320F (p value = 0.0027 and 0.0023, respectively), but not for mini-Gαs/i-Y320E (p value = 0.22; **Table S5**), indicative of mini-Gαs/i-Y320E being most compromised and as a result, not conforming to one site saturable binding model. Titration of acceptor expression levels was verified by flow cytometry during each repeat of the experiment (**Supplementary Figure 5d-f**).

These findings demonstrate that Y^320^ is a key determinant of GPCR coupling. We thus conclude that the corresponding phosphoevent at Gαi Y^320^ (which we observed in growth-factor treated cells) may be sufficient to uncouple the G protein from Gi-PCR, which is in keeping with prior observation that growth factor stimulation leads to the release of Gαi from CXCR4 (71). Because Y^320^ engages in (distinct) intramolecular H-bonds in both inactive and GiPCR-activated conformation (**Figure 4a**), the intermediate performance of the Y320F mutant in these receptor recruitment assays is not unexpected.

### Impact of the phosphosites on canonical GPCR-stimulated chemotaxis

Next we asked if the overall theme of phosphotyrosine-mediated inhibition that we observe in GPCR coupling (Y^320^), or the canonical Gαi-mediated cAMP (Y^320^) and Gβγ signaling (Y^154/155^ and Y^320^) cascades is physiologically relevant and translates into phenotypic changes in cells. To this end, we assessed the impact of these three tyrosines on directional migration, a.k.a, chemotaxis, because the CXCL12/CXCR4 system is known for its ability to support chemotaxis by triggering a Gαi-dependent signaling axis (52). We used a well-accepted transwell assay in which HeLa cells were allowed to migrate across a 0-40 nM CXCL12 gradient for 20 h in the presence of PTX (**Figure 5a**), a duration that was chosen based on prior work (60). The positive and negative controls in our assay behaved as expected, i.e., cells expressing the PTX-sensitive WTs G protein (negative control) did not migrate, whereas those expressing the PTX-resistant WTr G protein (positive control) migrated (**Figure 5b***-top panels*; **Figure 5c**). When compared against the PTX-resistant WTr G protein, all but one mutant showed various degrees of impairment in their ability to drive chemotaxis (**Figure 5b-c**); the YYY154-155- 320EEF mutant performed just as well as WTr. The YYY154-155-320EEF mutant was significantly more competent in chemotaxis in head-to-head comparisons with the virtually immotile Y320E, YYY154-155-320EEE and YY154-155EE mutants (*p* values 0.0002, 0.0017 and 0.004, respectively; **Table S6**). The Y320F mutant was significantly better at supporting chemotaxis compared to the Y320E mutant (*p* value 0.0289; **Table S6**). These findings indicate that the Y320E substitution was sufficient to inhibit chemotaxis, and the Y320F substitution was sufficient to reverse the inhibitory effects observed in Tyr/Y→Glu/E mutants within the interdomain cleft.

**Figure 5:**
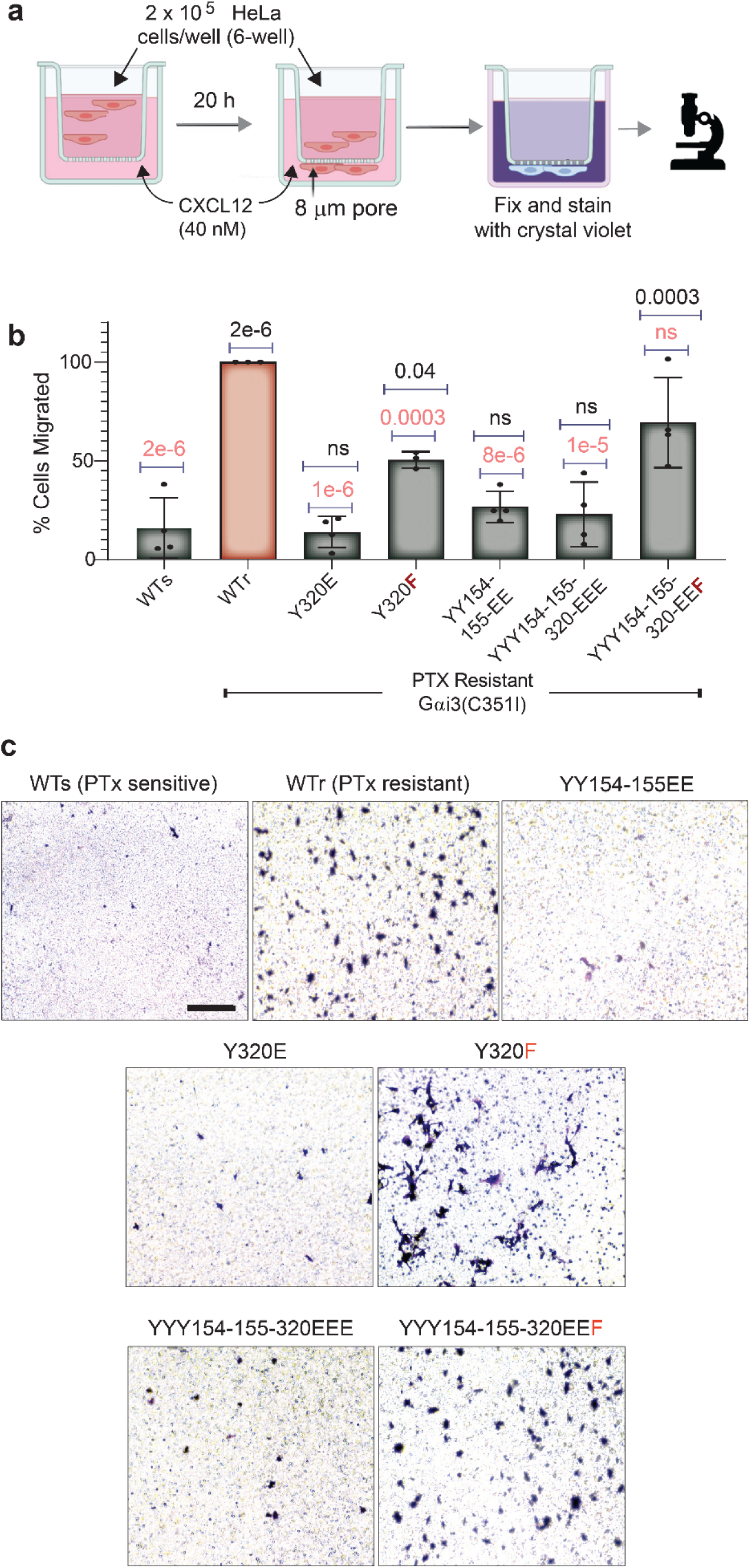
A non-phosphorylatable mutant of Y^320^ rescues CXCL12- stimulated chemotaxis. a. Schematic displays the pertinent details of a transwell cell migration assay to measure the chemotactic capability of HeLa cells transiently expressing various Gαi3 mutants across a 0 nM (top) to 40 nM (bottom chamber) CXCL12 gradient (see *Methods*). b. Bar graph shows % cell migration compared to cells expressing the wild-type PTX-resistant (WTr positive control) Gαi3. c. Representative images of crystal violet-stained transwell membrane show the cells that have migrated successfully towards the CXCL12 gradient. Scale bar: 100 µm. *Statistics*: *P* values were determined by one-way ANOVA followed by Tukey’s multiple comparison test. P-Values in black and red fonts indicate significance compared to the negative control wild-type sensitive (WTs) and the positive control resistant (WTr) G proteins, respectively. Error bars represent ± S.E.M; n=3-4 independent experiments, each with 3 technical replicates.

Taken together, our results indicate that Gαi phosphomodifications that we revealed in growth factor-stimulated cells generally inhibit the canonical GPCR-driven chemotaxis process, as exemplified by almost all the mutants, i.e., S/T phosphorylation at Ser^44^, Ser^47^ and Thr^48^ in the P-loop and tyrosine phosphorylation at the interdomain cleft and at the C-terminus. Consistent with all other readouts, the phosphomimicking mutation at Y^320^ was sufficient to inhibit chemotaxis, alone or in combination with YY154-155EE (**Table 3***-column #5*). Intriguingly, as in the Gβγ release and activation assay, making Y^320^ non-phosphorylatable (Y320F) rescued the deficiencies of not only the triple phosphomimic YYY154-155-320EEE but also the double mutant YY154-155EE.

### Structural analysis of phosphomimicking and non-phosphorylatable Gαi mutations

To put these experimental findings in a structural context, we projected the mutated residues on the Gαi structure in three structurally characterized functional states: inactive GDP-bound with Gβγ, nucleotide-free in complex with a GPCR and Gβγ (this transient state corresponds to nucleotide release), and active GTP-bound (**Figure 6a**). This revealed the proximity and interaction of all three P-loop residues with the nucleotides, with Thr^48^ directly coordinating the first phosphate for both GDP and GTP, Ser^47^ the Mg2+ ion in the GTP-bound state, and Ser^44^ forming a hydrogen bonding network with K(Lys)^45^ and the second GDP/GTP phosphate. These observations suggest that phosphorylation (or phosphomimicking mutations) at these positions would disrupt the ability of Gαi to bind nucleotides and hence participate in the GDP/GTP exchange cycle in response to GiPCR stimulation. In the nucleotide-free GPCR-bound state, these residues do not form any critical contacts which is likely why their mutations may trap this state, similar to dominant negative Gαi (66).

**Figure 6:**
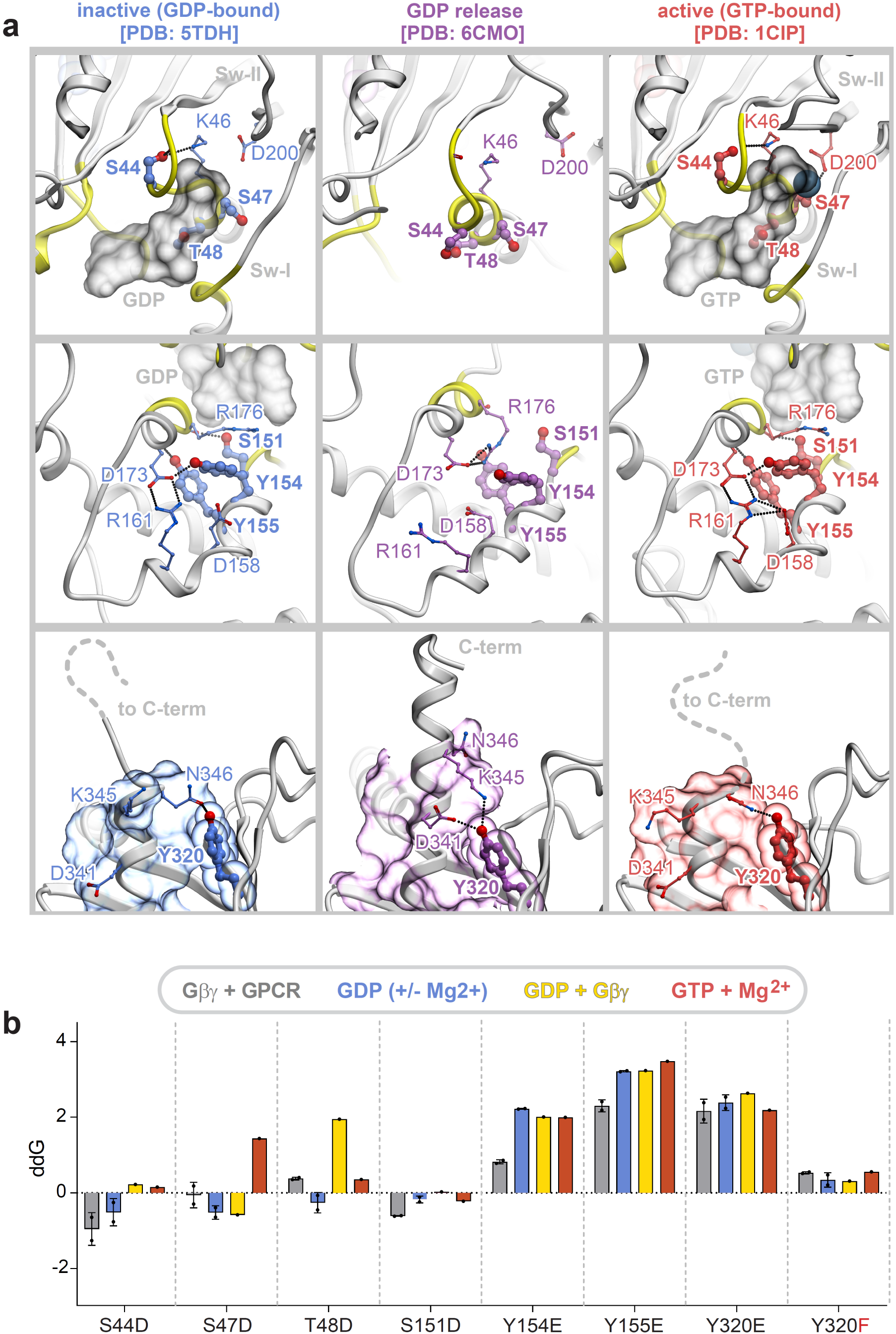
Computational prediction of the impact of residue mutation on the stability of Gɑi in various complexes. a. Crystal structure of WT Gαi3, highlighting contact of the residues in the P-loop region (*<u>Top panels</u>*: Ser^44^, Ser^47^ and Thr^48^), interdomain cleft (*<u>Middle</u>*: Y^154^ and Y^155^) and the C terminal α5 helix region packed against β4-β6 sheets (*<u>Bottom panels</u>*: Y^320^), in their GDP and Gβγ-bound inactive state (PDB: TDH), GDP release state (PDB: 6CMO) and GTP-bound active state (PDB:1CIP). H-bonds between the residues which were used in this study and its neighboring residues in the non-phosphorylated state. b. Bar graph displays the computationally predicted effects of phosphomimicking or non-phosphorylatable mutations on the stability of Gαi3 in various complexes and conformations, including a complex with Gβγ+GPCR (PDB:6cmo,6ot0), GDP (+/-Mg^2+^; PDB:1bof, 1gdd), GDP+Gβγ (PDB:1gg2), GTP+Mg^2+^ (PDB:1gia). Bar height indicates delta delta G [ddG], i.e., the predicted change in the change in Gibbs free energy of folding; higher / more positive bars reflect a larger degree of destabilization by the respective mutation while negative numbers indicate the opposite. See **Supplementary** Figure 6a for the relative stability of the different mutant state conformations and complex compositions compared to the Gβγ-bound state, and **Supplementary** Figure 6b for the extended analysis of both non-phosphorylatable substitutions (Ala, Phe) as well as phosphomimicking substitutions at each site.

The Ser^151^/Y^154^/Y^155^ residue cluster in the αE helix is engaged in a dense hydrogen-bonding network in both nucleotide-binding states, which is partially released in the domain-open, nucleotide-free state (**Figure 6a**). In fact, Y^155^ is fully buried in all structurally characterized states of Gαi, suggesting that for it to be phosphorylated the AHD must partially unfold. Y^154^, however, may become exposed through a rotamer change, especially in the nucleotide-free state.

The most interesting structural observations are related to Y^320^ in the β6 strand. In the nucleotide-bound states, this residue packs against the receptor-engaging C-terminal α5 helix and hydrogen-bonds to N346 (**Figure 6a**). In the process of Gαi activation by the receptor, due to a rotation and an upward shift of the α5 helix, this residue switches its hydrogen bonding to K^345^ and D(Asp)^341^, while continuing to provide the stabilizing/packing interactions for that helix. In other words, Y^320^ works both as a stabilizing residue and a conformational switch in the process of Gαi activation by a GPCR. We hypothesize that Y^320^ phosphorylation or its replacement by E(Glu) impairs the stability of the C-terminal α5 helix and may, for example, make it more prone for leaving its location in the center of the Gαi GTPase domain β-barrel (72, 73). By contrast, a Y320F substitution still allows this residue to serve its stabilizing role for the α5 helix, as only the hydrogen bond switching functionality is eliminated by that mutation.

Next, we computationally assessed the stability of the various structurally characterized functional states of Gαi when mutated at the phosphorylated positions. Four conformational states were assessed: nucleotide-free in complex with a GPCR and Gβγ, GDP-bound, GTP-bound with Gβγ, and GTP-bound), and multiple experimental structures were included where available. The effects of mutations on stability of each state was calculated and expressed as predicted ΔΔG of folding, with positive numbers indicating destabilization (state disfavored) and negative numbers indicating stabilization (state favored) (**Figure 6b** and **Supplementary Figure 6**).

S44D and S151D were predicted to slightly favor the nucleotide-free receptor- and Gβγ-bound state. In keeping with prior experiments (41, 66) S47D strongly disfavors the GTP-bound state of Gαi and T48D the GDP- bound state. Phosphomimicking mutations at Y154 and Y155 were predicted to be strongly destabilizing across the board, with a somewhat milder impact on nucleotide-free receptor- and Gβγ-bound Gαi. Similarly, Y320E was strongly destabilizing, with the GDP/Gβγ-bound state of Gαi affected the most (**Supplementary Figure 6**). By contrast, Y320F had only a minor destabilizing effect, and the GDP-bound states were affected the least (**Figure 6b** and **Supplementary Figure 6**).

In summary, these structural and computational observations provide a rational basis for the experimentally observed effects of the mutations.

### Impact of the phosphomimicking and non-phosphorylatable mutations on membrane localization of Gαi

The contrasting effects of the phosphomimicking Y320E (and YYY154-155-320EEE) and the non-phosphorylatable Y320F (and YYY154-155-320EEF) substitutions across all readouts (**Table 3***-columns #3-7*) led us to hypothesize that the non-phosphorylatable Y320F substitution may provide an advantage that drives the observed proficiency of the G protein in most assays. Because all mutants expressed at similar levels in cells (**Figure 2c-d**), difference in protein stability was deemed as an unlikely factor, and instead, we investigated protein localization. Confocal immunofluorescence studies carried out in HeLa cells revealed striking difference in the localization pattern between the E(Glu) and F(Phe) mutants (**Figure 7a**) Gαi proteins in which a Tyr(Y) or a Phe(F) occupy the position 320 (Y320F, YY154-155EE and YYY154-155-320EEF) localize predominantly at the peripheral membrane, presumably the plasma membrane (PM); however, proteins with a Glu/E at that position (Y320E and YYY154-155-320EEE) localize predominantly in the cytosol. Furthermore, cell fractionation assays confirmed these findings; Gαi proteins with an intact Y^320^ (WTr or YY154-155EE mutant) or in which the Y^320^ was substituted with Phe(F) (EEF and Y320F mutants) were detected in the crude membrane fractions, indistinguishable from each other. However, the Gαi proteins in which the Y^320^ was substituted with Glu(E) (Y320E and EEE mutants) were detected primarily in cytosolic fractions (**Figure 7b, c**).

**Figure 7:**
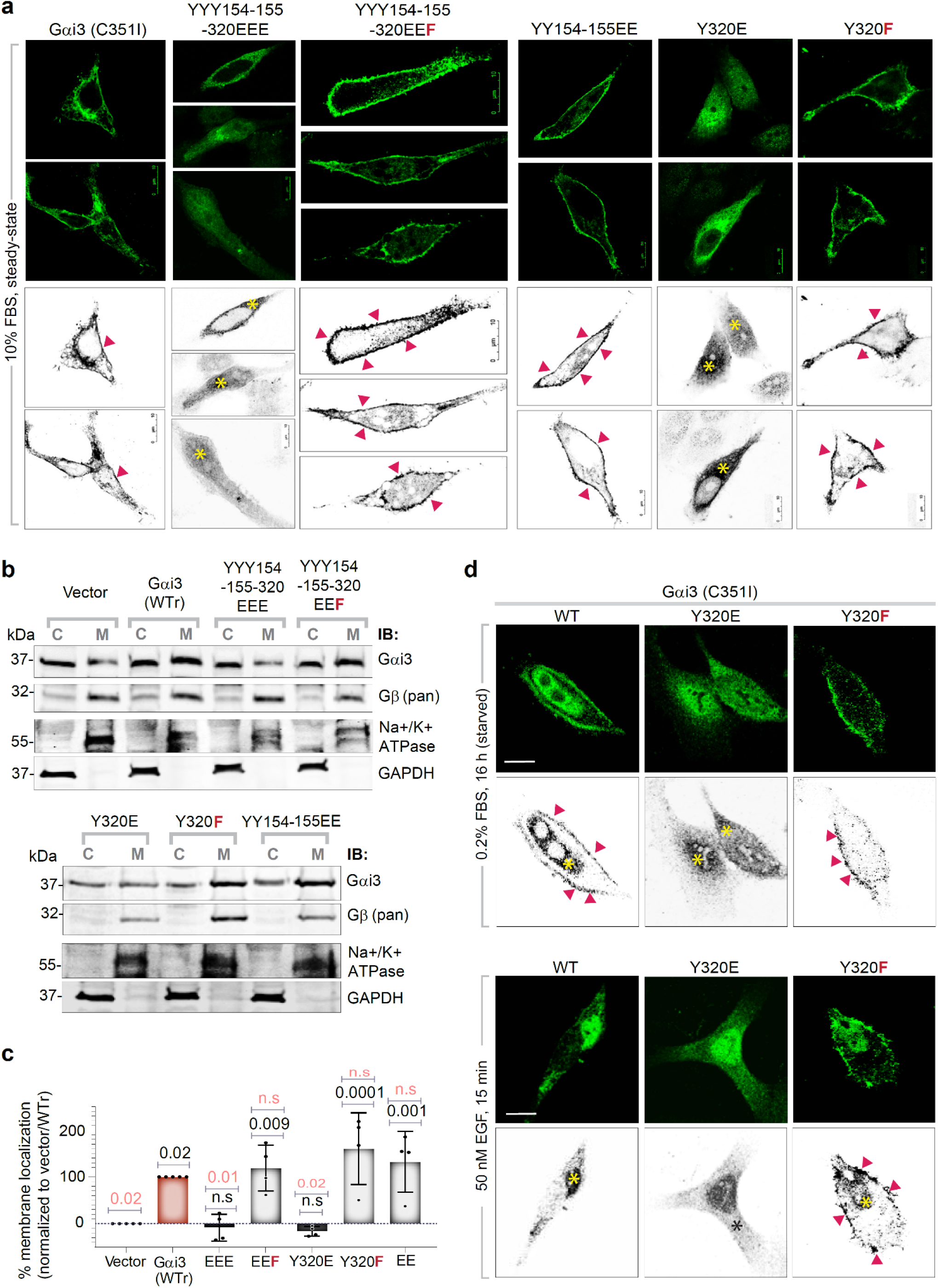
Effect of phosphomimicking or non-phosphorylatable Gαi3 mutants on protein localization. a. The subcellular localization of the indicated Gαi3 constructs was assessed in transiently expressing HeLa cells by confocal immunofluorescence microscopy. Representative images are displayed as assembled montage in panel a. *<u>Upper panels</u>*: green, Gαi3). *<u>Lower panels</u>*: green channels depicting Gαi3 alone are presented in grayscale (red arrowheads indicating peripheral membrane localization; yellow asterisks indicating intracellular localization). Scale bars: 10 µm. b. Membrane localization of the indicated Gαi3 constructs and pan-Gβ were assessed by subcellular fractionation of post-nuclear supernatants derived from the homogenates of each transiently expressing HeLa cell line in panel a. Immunoblots representative of 3 independent biological repeats are shown. C, cytosol; M, crude membranes. Cytosol and membrane fractionations were assessed using the markers Glyceraldehyde-3- phosphate dehydrogenase (GAPDH) and β1 Sodium Potassium ATPase (ATP1B1), respectively. c. Immunoblots in b were quantified by band densitometry and displayed as bar graphs. Results are expressed as % distribution of Gαi3 on membrane fractions on a set scale of 0-100%, where 0% is the vector alone (endogenous Gαi3; negative control) and 100% is the PTX-resistant Gαi3 (WTr, positive control). *P* values were determined by one-way ANOVA with Tukey’s multiple comparison’s test. P-Values in black and red fonts indicate significance compared to the negative control (vector) and positive control (WTr) conditions, respectively. Error bars represent ± S.E.M; n=4 independent experiments. d. The subcellular localization of the indicated Gαi3 constructs was assessed in transiently expressing HeLa cells and treated with and without EGF (50nM) for 15 mins and visualized by confocal immunofluorescence microscopy. Representative images are displayed as assembled montage in panel a. Upper panels: green, Gαi3). Lower panels: green channels depicting Gαi3 alone are presented in grayscale (red arrowheads indicating peripheral membrane localization; asterisks indicate intracellular localization). Scale bars: 10 µm.

These findings demonstrate that Y^320^ on Gαi is a key determinant of G protein localization. That the Y320E mutant localizes to the cytosol but Y320F localizes at the PM, suggested that phosphorylation at Y^320^ upon EGF stimulation may be sufficient for Gαi mislocalization. To test if such is the case, we assessed protein localization by confocal microscopy in cells with or without EGF stimulation. In serum starved cells, WT Gαi localized to the PM (red arrowheads; **Figure 7-***left*) and in the perinuclear region, presumably at the Golgi (as shown previously (74); yellow asterisk; **Figure 7d-***left*); upon EGF stimulation, the PM localized pool was selectively lost. Such ligand-dependent changes in PM localization were abolished in the Y320E/F mutants; PM localization was undetectable in the case of the constitutive phosphomimic Y320E mutant (**Figure 7d-***middle*) and stably detected in the non-phosphorylatable Y320F mutant (red arrowheads; **Figure 7d-***right*) regardless of ligand stimulation. Furthermore, that the YY154-155EE and YYY154-155-320EEF mutants localize at the PM but YYY154-155-320EEE localizes to the cytosol indicates that phosphorylation at Y^320^ may be required for Gαi to lose PM localization (**Table 3***-column #8*).

## Discussion

Canonical trimeric G protein signaling is triggered by ligand activated GPCRs; however, the canonical GPCR pathways have been shown in many contexts to be subject to crosstalk and regulation by growth factors. Here we reveal the molecular mechanisms that enable growth factor RTKs to modulate the canonical G protein signaling. More specifically, we show that growth factors may shape canonical signaling, in part, through phosphorylation of Gαi (**Figure 8**). Phosphorylation occurs at key residues (clusters of residues) located at three strategic hot spots within the GTPase: the P-loop, the interdomain cleft and the C-terminus. Using a library of phosphomimicking mutants in a series of assays to interrogate some of the most upstream (receptor recruitment and trimer dissociation) as well as downstream (cAMP inhibition and chemotaxis) events in canonical Gi/GPCR signaling, we show that the phosphomodifications predominantly inhibit ligand-induced signaling with some of them also promoting constitutive Gβγ signaling. These findings illuminate the basis for some poorly-understood observations by others that growth factors (EGF and IGF1) inhibit Gi coupling to and activation by GPCRs (CXCR4) (30, 50, 71). Because phosphorylation of Gαi at Y(Tyr)^154^ and Y(Tyr)^155^ by RTKs such as EGFR increases its basal GDP-to-GTP exchange rate by ∼15.6-fold (5), the current work implies that phosphomodulation of Gαi downstream of growth factors may serve as a key determinant of whether a G protein may be activated by canonical vs. non-canonical GPCR-independent mechanisms. While physiologic growth factor signaling may spare the G protein for canonical signaling downstream of GPCRs, overzealous signaling in pathologic conditions such as cancers may steal the G protein from canonical GiPCR-triggered pathways (this work), and instead, facilitate the transduction of non-canonical signals (shown earlier (5)). Such a steal may facilitate contextualization of signals; for example, although phosphorylation of the interdomain tyrosines (Y^154^Y^155^) could suppress cAMP upon growth factor stimulation and enhance migration during wound closure or invasion through matrix proteins (5), such phosphorylation (mimicked by YY154-155EE in this work) was unable to suppress cAMP or support chemotaxis upon GiPCR stimulation. We also show that the mechanism(s) for inhibition of canonical signal differs based on the location of the phosphosites within the G protein (see **Figure 8**; **Table 3**-*columns #9-10*). These findings are discussed below.

**Figure 8.**
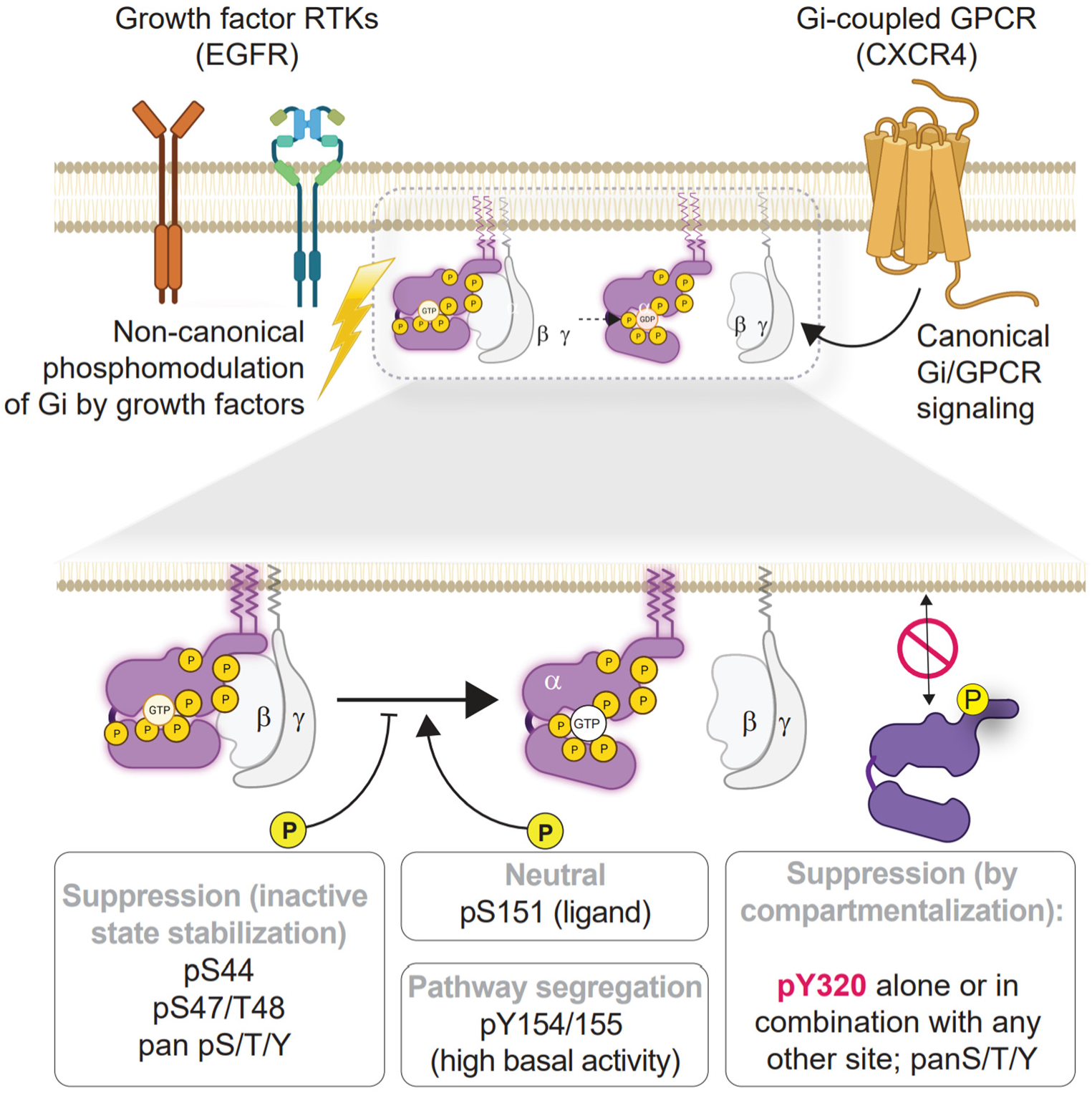
Schematic of the major findings in this work. Growth factors may have at least 3 distinct effects on canonical Gα(i)βγ/GPCR signaling via phosphorylation of Gαi. Phosphorylation of some clusters of residues may suppress signaling by stabilizing the GDP-bound conformation (S44) or stabilizing the GDP/Gβγ-bound state (S47/T48) and thereby impeding ligand-stimulated nucleotide exchange and trimer dissociation (S44, S47, and T48). Phosphorylation at other sites may either have no effect (S151) or result in a segregation (Y154, Y155) between non-canonical (growth factor RTK→Gi) and canonical (Gi/GPCR) pathways, while maintaining high basal activity due to defective Gβγ sequestration. Finally, phosphorylation at one key site may compartmentalize signaling via the regulation of G protein localization at the PM. See also **Table 3** for a comprehensive summary of all mutants tested in this work.

First, phosphorylation of key residues at the interdomain cleft, as Ser^151^, Y(Tyr)^154^ and Y(Tyr)^155^ (which are phosphorylated in combinations of pSer^151^ alone and pY^154^/pY^155^) we find that while Ser^151^ has little or no impact, the pair of Ys impair ligand stimulated Gβγ release, partially impair ligand stimulated cAMP suppression, and inhibit chemotaxis. Importantly, all of this occurs in the background of elevated constitutive Gβγ signaling due to the impairment of the mutated/phosphorylated Gαi ability to bind and sequester/inactivate Gβγ. Computational analyses revealed that phosphorylation at Y^154^/Y^155^ may favor the nucleotide-free conformation. These findings are consistent with our prior work (using non-phosphorylatable mutants in FRET-based assays) which showed that dual phosphorylation at Y^154^/Y^155^ is required for growth factor stimulated trimer dissociation and increases basal nucleotide exchange rate (∼5-fold higher than WT) (5). Increased basal activity effect was attributed to disrupted intermolecular interactions and packing (loss of H-bonds) within the inter-domain cleft, which may affect opening/closing of the protein’s all-helical and Ras-like domains (5). Although molecular dynamic simulation studies have shown that domain opening is insufficient for GDP release (75), phosphorylation at Y^154^/Y^155^ may affect neighboring residues within the nucleotide-binding pocket, such as those in the αD-αE loop which includes the so-called ‘NDS^151^’ motif. Alternatively, because both Y^154^ and Y^155^ face towards αF and Sw-I, particularly Y^155^, it is possible that destabilization of Sw-I could serve as a mechanism for pTyr-induced allosteric activation of the GTPase; Sw-I was recently identified as a key conduit in the allosteric path to nucleotide release that is triggered by the guanine-nucleotide exchange factor, GIV/Girdin, transmitting forces from the Sw-II to the hydrophobic core of the GTPase (76). While the identity of the kinase that phosphorylates S^151^ remains unknown, Y^154^/Y^155^ has been shown to be phosphorylated by multiple RTKs (but not non-RTKs) (5). Regardless of who phosphorylates them and how these phosphosites impact the basal exchange rate, what is clear is that they functionally uncouple the G protein from GPCRs, by reducing Gβγ-binding and trimer assembly.

In doing so, the interdomain cleft events segregate the RTK→Gαi pathway (with resultant higher basal activity) from the canonical GPCR→Gαi pathway.

Second, phosphorylation of key residues within the P-loop, i.e., Ser^44^, Ser^47^ and Thr^48^ (the latter two are simultaneously phosphorylated) impairs ligand-stimulated Gβγ release, and cAMP suppression, but largely spares the ability of the G protein to bind Gβγ and assemble trimers. Computational analyses revealed that phosphorylation at Ser^47^ preferentially destabilizes the GTP-bound state, whereas phosphorylation at Thr^48^ destabilizes the GDP-Gβγ-bound Gαi. These are not unexpected, because the P-loop, which is essentially the sequence ^40^GAGESGKST^48^ (77), is known to participate in nucleotide binding and coordination of Mg^2+^ (78); while Ser^47^ coordinates the Mg^2+^, T48 bridges the β and γ phosphates in GTP. Our findings are consistent with prior work in which phosphorylation at Ser^44^ on Gαi2 was implicated as a basis for desensitization of μ-opioid receptors (79). Similarly, the S47R mutant Gαi3 [implicated in auriculo-condylar syndrome (ACS)] produces dominant-negative proteins that couple to the GPCR but cannot bind and hydrolyze GTP (80). A similar effect has been observed in the case of Gαq (Ser^53^, which corresponds to Ser^47^ in Gαi); phosphorylation of this site impedes GTP-binding and downstream signaling events (81). Although the identity of the kinase(s) that phosphorylate these P-loop sites remains unknown, the phosphorylation functionally uncouples the G protein from GPCRs, by reducing ligand-induced Gβγ-release but without elevating the constitutive Gβγ activity. Because coupling to Gβγ and GPCR is unlikely to be impacted, Gαi proteins phosphorylated at these sites may behave as dominant negative proteins that couple to the GPCR but cannot release Gβγ; nor will they bind and hydrolyze GTP. Thus, the P-loop sites may not only inhibit the canonical GPCR→Gαi pathway but carry the potential to exert a dominant negative effect.

Third, phosphorylation of Y^320^, a key residue at the C-term β6 strand, impairs Gβγ-binding and ligand-stimulated Gβγ release, receptor coupling, ligand stimulated cAMP suppression, and inhibits chemotaxis. A prior study had established the importance of this residue in coupling to GPCRs (41), and we previously established that Y^320^ is phosphorylated by RTKs, both in vitro and in cells (5). We found that phosphomimicking mutants of Y^320^, either alone or in combination with other tyrosines in the interdomain cleft, renders the G protein inert to canonical signals. Interestingly, a Y320F substitution not just retains Gβγ-binding, but also ligand-stimulated Gβγ release. This is consistent with the nature of this residue being a Phe in selected other subtypes of Gα proteins, e.g. in Gα15 and Gα12 (**Supplementary Figure 2**). To our surprise, Y320F could reverse defects in signaling observed in the proteins in which the two Y(Tyr) in the interdomain cleft region is phosphorylated; we believe that this effect is primarily via restoring the ability of Gαi to bind Gβγ and assemble trimers (see **Table 3**). Y320F also reversed the effects of Y320E in pan-Tyr mutants; we believe that this is because of the contrasting patterns of protein localization—proteins with Y320E (alone or in combination with mutants of the interdomain cleft) were cytosolic, whereas proteins with Y320/Y320F (alone or in combination) were localized to the PM. As for the mechanism by which Y^320^ dictates membrane localization, it is unlikely that the mechanism is either of the two known determinants of membrane localization of Gαi subunits, that is, N-terminal myristoylation and/or palmitoylation and association with Gβγ dimers (17) (YY154-155EE, which is deficient in Gβγ-binding, is localized at the PM). What was also puzzling is that Y320E alone was sufficient to abrogate Gβγ-binding. Recently revealed structural insights (73) into how the G protein chaperone and guanine nucleotide exchange factor Ric- 8 binds Gαi sheds some light on this puzzle. The core of the Gαi•Ric8 complex requires the β4-β6 strands of Gαi and Y^320^(β6) serves as a contact site (73), which raises the possibility that it may be a critical determinant of the interaction. It is noteworthy that the Y320E containing Gαi mutants in our system mimic the defects previously reported in Ric8-depleted diverse systems (fungi, fly, worm, and mammalian), i.e., Gαi subunits show reduced steady-state abundance and localization at the PM, associate inefficiently with Gβγ dimers and consequently, assemble heterotrimers poorly (72, 82–86). Whether phosphorylation of Gαi at Y^320^ impacts its interaction with and modulation by Ric8 remains to be explored. Regardless, what emerged from this work is that phosphorylation at Y^320^ by growth factor RTKs impairs GPCR coupling and PM localization (cytosolic compartmentalization) of the Gαi protein and thereby inhibits the canonical GPCR→Gαi pathway.

Finally, it is noteworthy that besides Gαi, GIV also modulates Gαs (37), raising a possibility that GPCRs coupled to Gαs may also be similarly impacted via phosphomodulation by EGFR. We noted that while Ser^44^, Ser^47^, Thr^48^ and Y^154^ are conserved in Gαs, Ser^151^ and Y^155^ are not (**Supplementary Figure 2**). Although Y^320^ is conserved in Gαs, the flanking sequence is not, suggesting that RTK-based recognition of substrate site may not happen. Consistent with this, we previously showed that Gαi, but not Gαs is a preferential substrate of EGFR (5). By contrast, the non-RTK Src phosphorylated all G proteins tested equally efficiently, including Gαs and Gαi (5).

As for study limitations, we did not evaluate the myriad of downstream signaling pathways that are triggered by CXCL12; their dissection is a must for us to assess the full impact of the phosphomodifications. Neither did we reveal the identity of the Ser/Thr kinases that phosphorylate the G protein. It is possible that the phosphoevents we report here exert their effects either independently of each other or influence each other (perhaps resulting in some logical sequence and/or hierarchy). Although these aspects remain unknown, any phosphosite which accelerates nucleotide exchange can be expected to enhance the phosphorylation of residues that are buried within the interdomain cleft and hence, are otherwise inaccessible to kinases. Although the fundamental nature of our findings are expected to be widely relevant across cell types, it is possible that our findings in HeLa cells may be uniquely impacted by the endogenous abundance of Gβγ subunits and AC isoforms.

In conclusion, we show how diverse Gαi phosphoevents that are observed in growth factor-stimulated cells exert a coherent effect, i.e., they all inhibit canonical GPCR signaling via distinct mechanisms that are dictated by the location of the phosphosites. Findings provide mechanistic into how growth factors inhibit canonical Gi/GPCR signaling insights from the level of the atoms to the level of the whole cell.

## Materials and Methods

### Regents and consumables

All the regents and consumables and software programs used in this work is listed (with identifiers) in a key resource table in Supplementary Materials.

### Constructs and cloning

Wild-type rat Gαi3 cDNA, obtained from A. Spiegel (National Institutes of Health, Bethesda, MD) was subcloned into pcDNA 3.1 and validated previously (90) . Using this construct as template, we created all other Gαi3 mutants featured in this work using a QuikChange® Site-Directed Mutagenesis Kit as per manufacturer’s protocol (Stratagene, USA). A variant that is insensitive to ADP-ribosylation by Pertussis Bordetella toxin (PTX) (45) was created for each construct by mutating residue cysteine (Cys; C) 351 into Isoleucine (Ile; I) (93).

A BRET-based CAMYEL (cAMP sensor using YFP-Epac-RLuc (68)) sensor was a gift from Paul Insel and Tracy Handel (University of California, San Diego). V2-CAMYEL, a modified version of the sensor with increased luminescence was generated by site-directed mutagenesis of the RLuc region (C124A/M185V) (88).

Recombinant CXCL12 was expressed and purified exactly as done before (94). mVenus-hGBB1 and mVenus-hGBG2 were a gift from Nevin Lambert, Univ. of Alberta (89). Mini-Gαi construct was a gift from Nevin Lambert, Augusta University, Augusta, Georgia, USA. The mini-Gαs/i is a chimaera of Gαs and Gαi, keeping in parity the Y^320^ residue of Gαi3 and Gαs the Y320E and Y320F mutants were generated using a QuikChange® Site-Directed Mutagenesis Kit as per manufacturer’s protocol (Stratagene, USA).

### Pulldown assays

These studies were carried out exactly as described previously (36, 91, 92, 95). For in vitro pull-down assays, equimolar amounts of bacterially expressed and purified GST-tagged proteins or GST alone (negative control) immobilized on glutathione-Sepharose beads were incubated with transfected with required constructs and pre-cleared lysates of Cos7 cells in binding buffer [25 mM Hepes (pH 7.4), 100 mM NaCl, 10 mM MgCl_2_, 30 μM GDP, 2 mM DTT, 0.5% Triton X-100, 1x Complete Protease Inhibitor Mixture (Roche), and 1× Phosphatase Inhibitor Mixtures 1 and 2 (Sigma)]. Bead-bound proteins and lysates were incubated with constant tumbling at 4 °C for 4 h. Beads were washed four times with 1 mL of lysis buffer and were eluted in 1× Laemmli sample buffer by heating to 95 °C for 5 min and analyzed by SDS-PAGE. Bound total and phosphorylated Gαi3 proteins was analyzed by dual color immunoblotting. In the case of His-GIV-CTs pulldown studies, the protein was bound to HisPur™ Ni-NTA Resin (Thermo Fisher) and the remaining steps were carried out as for pulldown assays with GST proteins.

### In cellulo phosphorylation of Gαi3 protein and linear-ion-trap mass spectrometry

Cos-7 cells were cultured according to American Type Culture Collection (ATCC) guidelines. For *in vivo* phosphorylation assays on Gαi3, Cos-7 cells were transiently transfected with wild-type Gαi3 C-terminally tagged with FLAG, and serum starved for 16 h (0 % FBS) prior to stimulation with EGF (50 nM, 15 min) in the presence of the cell-permeable tyrosine phosphatase inhibitor, sodium orthovanadate (100 μM, added 1 hr prior to EGF stimulation). Reactions were stopped using PBS chilled to 4°C and supplemented with 200 μM sodium orthovanadate and Ser/Thr phosphatase inhibitor (Sigma), after which cells were immediately scraped and lysed. Cell lysates for mass spectrometry were prepared by resuspending cells in lysis buffer [20 mM HEPES, pH 7.2, 5 mM Mg-acetate, 125 mM K-acetate, 0.4% Triton X-100, 1 mM DTT, supplemented with sodium orthovanadate (500 µM), phosphatase (Sigma) and protease (Roche) inhibitor cocktails], after which they were passed through a 28G needle at 4 °C, and cleared (10,000 x g for 10 min) before use in subsequent experiments. Equal aliquots of lysates were used as starting material for immunoprecipitation of Gαi with an antibody that recognizes FLAG (Sigma) and protein G beads.

To determine *in vivo* phosphorylation states of the FLAG-Gαi3 we used the QTRAP 5500 in the selected reaction monitoring (SRM) mode to scan for all possible phospho-forms of this protein. For this purpose, the bead-antibody-protein complex was trypsinized in solution with the aid of Rapigest© surfactant and analyzed using six distinct tandem mass spectrometry acquisition methods using a quadrupole time of flight (QSTAR Elite ABSCIEX) instrument and the ABSCIEX QTRAP 5500 (hybrid mass spectrometer). The acquisition methods were: 1) Data dependent MS2 method using the QSTAR Elite, 2) Data dependent MS2 method using QTRAP 5500, 3) Multiple reaction monitoring (MRM) methods for all possible phospho-forms of target protein using the QTRAP 5500 [**Supplementary Data File 1**], 5) MRM triggered full scan MS2 methods for the detected peptide phosphoforms using the QTRAP 5500 in the trap mode, 5&6) Precursor ion scans for -79 (in negative ion mode for S/T[P]) and 216.04 (in positive ion mode for Y[P]) ions. Upon detection of these indicator ions the mass spectrometer switches to ion trap mode to collect MS2 spectrum. SRM methods were developed for all possible tryptic peptides in phosphorylated and non-phosphorylated states. The ABSCIEX SRM Pilot^TM^ software was used for SRM method development. Ultimately a method with 210 SRM transitions states was developed for phosphorylated and non-phosphorylated tryptic peptides of Gαi3. In most cases there were at least 2 transitional states used for a given peptide mass. A total of 13 unique phosphorylation sites in the Gαi3 protein were detected by the QTRAP 5500, of which 3 were tyrosines [**Supplementary Data File 2**]. Because samples were not subjected to phosphoenrichment prior to Mass Spectrometry analyses, stoichiometry of any phosphoevent was calculated based on the phosphorylated over total peptides of any given sequence [**Supplementary Data File 1**].

We used another 10 μL of the same tryptic sample used in the previous SRM experiment, to run the QTRAP 5500 mass spectrometer in the “precursor ion scanning mode” either for an ion at m/z 79 in negative ion mode for serine and threonine phosphorylation, or an ion at m/z 216.043 for tyrosine phosphorylation in the positive ion mode. Once the precursor ions are detected, the instrument switches to positive ion trap scanning mode to isolate the parent ions and to carry out MS2 analysis on these ions. The collected MS2 spectra were analyzed using the ProteinPilot® search engine to identify the matching protein sequence from a database. The phosphorylation status of residues of the Gαi3 protein were traced and the ones we used are listed in **Table 1**.

### Sequence alignment construction

For **Supplementary Figure 2**, the amino-acid sequences of human, mouse, and rat Gα proteins were downloaded from UniProt (96) using the query ’family:“G-alpha family”’ and aligned the modified version of the S B Needleman-Wunsch method (97) as implemented in ICM (98), a software for Molecular Modeling (Biased Probability Monte Carlo and Molecular Dynamics) and Docking. The alignment was then projected to the amino-acid positions of interest (aa 44, 47, 48, 144, 151, 154, 155, 320) in rat Gαi.

### GiPCR-directed Gβγ activation assay

HeLa cells were cultured according to ATCC guidelines in DMEM supplemented with 10% FBS and an Antibiotic-Antimycotic solution (see Key Resource Table). On day 1, HeLa cells were plated at 800 K cells per well in a 6- well plate in DMEM supplemented with 10%FBS. On day 2, cell culture media was replaced by fresh DMEM supplemented with 10% FBS, after which cells were transfected with 0.5 µg/well VenusCT-Gβ, 0.5 µg/well VenusNT-Gγ, 1 µg/well of WT or mutant Gαi3, and 100 ng/well of GRK3tagged RLuc8 (total of approximately 2 µg/well of DNA) using TransIT-X2 transfection reagent according to manufacturer’s instruction. On day 3, cells were lifted from the wells in the 6-well plate using trypsin, transferred into 1.5 mL microcentrifuge tubes, spun down, and resuspended to 700 K cells/mL in DMEM (no phenol red) supplemented with 10% FBS. Equal aliquots (100 µL/well, which is ∼70 K cells per well) of the cell suspension were replated on a 96-well white/clear bottom plate and allowed to adhere for 5-6 h in a 37℃/5% CO_2_ incubator. After that, for PTX-treated conditions, cell media was carefully removed from the wells and replaced with 100 µL/well DMEM (no phenol red) supplemented with 10% FBS and 200 ng/mL PTX. On day 4, cell culture media was carefully removed from the wells and replaced by 80 µL/well of serum-free assay buffer (PBS with 0.1% D-Glucose and 0.05% BSA). The luciferase substrate Coelenterazine-h was added to each well (10 μL of 0.1 mM stock, for the final concentration of 10 μM). The plate was incubated at room temperature for 5 min, after which repeated readings of light emission at 485 and 515 nm were initiated using the Spark 20M plate reader (Tecan Lifesciences, Switzerland) and continued for 3 min for basal reads. Next, the plate was taken out of the luminometer, 10 µL of either buffer or 1 µM CXCL12 (for 100 nM final CXCL12 concentration) were added to each well, and the plate was read for additional 10 min, to assess agonist-stimulated Gβγ activation. BRET at each time point was expressed as intensity at 515 nm divided by intensity at 485 nm. For the generation of basal association bar graphs, results are integrated over the 3 min basal reads and normalized to the observed basal activity in the absence (pcDNA, set at 0%) or presence (PTX-sensitive WTs or and PTX-resistant WTr, set at 100%) exogenously transfected Gαi3. For ligand-induced association, the reads were first baseline-corrected by subtracting the average of basal reads, then integrated over 10 min post addition of agonist (12 reads), and then normalized to the range of 0% (PTX-sensitive Gαi3, WTs) to 100% (PTX-resistant Gαi3, WTr). The experiment was repeated in four independent biological replicates on different days, each containing three technical replicates. To increase statistical power, in the basal association experiments, PTX-treated and non-PTX treated samples from the same day were used as separate biological replicates thus producing n=8. The findings were displayed as graphs using GraphPad Prism 9. An average of three to four biological replicates is shown.

### CAMYEL Gαi-directed cAMP suppression assay

On day 1, HeLa cells were plated at 600 K cells per well in a 6 well plate in DMEM supplemented with 10% FBS. On the day 2, cell culture media was replaced by fresh DMEM supplemented with 10% FBS, after which cells were transfected with 0.5 µg/well WT or mutant Gαi, 1 µg/well of V2-CAMYEL (total 1.5 µg/well of DNA) using TransIT-X2 transfection reagent according to manufacturer’s instruction. On day 3, cells were lifted from the 6- well plate using trypsin, transferred into 1.5 mL microcentrifuge tubes, and resuspended in DMEM+10% FBS to 700 K cells/mL. 100 µL/well of the cell suspension (70 K cells per well) were replated on a 96-well white/clear bottom plate and allowed to adhere for 5-6 hours in a 37℃/5% CO_2_ incubator. After that, for PTX-treated conditions, cell media was carefully removed from the wells and replaced with 100 µL/well DMEM (no phenol red) supplemented with 10%FBS and 200 ng/mL PTX. Cells were cultured in the incubator overnight. On day 4, cell culture media was carefully removed from the wells and replaced by 70 µL/well of serum-free assay buffer (PBS with 0.1% D-Glucose and 0.05% BSA). The luciferase substrate Coelenterazine-h was added to each well (10 µL of a 0.1 mM stock, to achieve a final concentration of 10 µM), along with 10 µL of 1 mM stock of 3-isobutyl- 1-methylxanthine (IBMX), for the final concentration 100 μM IBMX. The plate was incubated at room temperature for 5 min, after which repeated readings of light emission at 485 and 515 nm were initiated using the SPARK 20M plate reader (Tecan, Switzerland) and continued for 3 min (for calculating the basal CAMYEL signal). Next, the plate was taken out of the luminometer and 10 µL of either buffer or 1 µM CXCL12 (for 100 nM final CXCL12 concentration) were added to each well. The luminescence 485 and 515 nm was immediately read for 6 min. Next, 10 μl of 100 μM forskolin (FSK; final in-well concentration 10 µM) were added, and the 485/515 nm luminescence was read for another 15 minutes. cAMP signal was calculated as the inverse BRET ratio (emission at 485 nm/emission at 515 nm).

For the generation of bar graphs, the average 1/BRET over 3-6 min preceding FSK addition was subtracted from the average 1/BRET over 10-12 min immediately post FSK addition. Results are expressed as fold change compared to the observed basal activity in the PTX-sensitive (WTs, set at 0%) and PTX-resistant (WTr, set at 100%) Gαi3 in the PTX-treated condition. The experiment was repeated in four independent biological replicates on different days, each containing three technical replicates. The findings were displayed as graphs using GraphPad Prism 9. In main-text figures, an average of three to four biological replicates is shown; in **Supplementary Figure 4b**, three separate biological replicates are shown.

### Recruitment of mini-Gαs/i to CXCR4-Rluc3 (acceptor titration)

For studying recruitment of G protein to ligand-activated GPCR, we used an established BRET based mini-Gαs/i recruitment assay (99). On day 1, HeLa cells were passaged and plated at 800 K cells per well in 6-well tissue culture plates in DMEM supplemented with 10% FBS. On day 2, cells were transfected with mixtures containing a fixed amount (100 ng/well) of HA-CXCR4-RLuc3 DNA (donor) and varying amounts (0, 156.25, 312.5, 625, 1250, 2500 ng/well) of mVenus-miniGαs/i WT, mVenus-miniGαs/i(Y320E) or mVenus-mini-Gαs/i(Y320F) DNA (acceptor); the total amount of DNA was complemented to 2.6 ug/well with pcDNA. Transfections were performed with TransIT-X2 transfection reagent according to the manufacturer’s protocol. On day 3 cells were lifted from the 6-well plate with trypsin, transferred into 1.5 mL microcentrifuge tubes, and resuspended in DMEM (without phenol red) supplemented with 10% FBS to 800 K cells/mL. Equal aliquots of cell suspension (100 μL/well; ∼80 K cells per well) were replated on a 96-well white/clear bottom plate and allowed to adhere in a 37℃/5% CO_2_ incubator. Additionally, for each transfected sample, 200 K cells per well were plated of a 12 well plate for flow cytometric analysis. On the day of assays, the media was replaced with serum free BRET assay buffer (PBS with 0.1% D-Glucose and 0.05% BSA), coelenterazine-h was added to the wells to the final concentration of 10 µM, the plate was incubated for 2 min, and basal reads (emission at 485 nm and emission at 515 nm) were collected for 5 min in a SPARK 20M plate reader (Tecan, Switzerland). After this, either buffer or CXCL12 was added to the plate, the latter to a final concentration of 100 nM, and readings were recorded for an additional 10 min. BRET ratio was calculated as emission at 485 nm divided by emission at 515 nm.

For the generation of bar graphs, BRET readings were baseline-corrected and control-subtracted (the former using reads preceding agonist addition, the latter reads from buffer-treated wells), and the average over 5-8 readings (approximately 15 min) after CXCL12 addition was calculated. The experiment was repeated in five independent experiments on different days, each containing four technical replicates. The findings were displayed as graphs using GraphPad Prism 9. An average of five biological replicates is shown. For model fitting and parameter comparison between data specific fit and global fit we used extra sum-of-squares F test as implemented in GraphPad Prism. For Lineweaver-Burk plots comparison we used F-test to compute the significance of non-zero slope, also implemented in GraphPad Prism.

### Flow cytometry

Cells transfected for the mini-Gαs/i recruitment assay above were seeded at ∼200K/well in a 12 well plate. The next day cells were rinsed with PBS, detached with Accutase, and lifted in DMEM supplemented with 10% FBS, after which 300µL of each cell suspension (containing ∼100K cells) was transferred into a conical bottom 96 well plate. The plate was centrifuged at 400g for 5 min, supernatant aspirated, and cells resuspended in 200 µL/well PBS with 0.5% BSA (flow buffer). This was done twice. After the final spin, cells were resuspended in 50 µL flow buffer with of 1:1000 dilution of APC-conjugated HA antibody in PBS supplemented with 0.5% BSA; no antibody was added to the control well. Cells were stained on ice in the dark with rocking for 30 min, then washed twice with 200 μL/well of flow buffer. Finally, cells were resuspended in 300 μL of flow buffer and data was collected in Guava EasyCyte benchtop flow cytometer (Luminex, USA). Data were analyzed in FlowJo (10). A representative biological replicate of the flow cytometry analysis is shown in **Supplementary Figure 5d-f**.

### Transwell Migration Assays

This assay is based on the Boyden Chamber method (100), which measures the chemotactic ability of cells through pores in a membrane insert toward a chemoattractant (in this case CXCL12). On day 1, 1M HeLa cells per well were plated in a 6-well plate in DMEM media supplemented with 10% FBS. On day 2, cells were transfected with 2 μg/well DNA of WTs, WTr and the other mutants. After 36 h, cells were treated with 100 ng/ml PTX for 12 h. On the following day, cells were rinsed in PBS, detached with trypsin, lifted in DMEM supplemented with 0.4% FBS, centrifuged, and resuspended in DMEM supplemented with 0.4% FBS. Cells were seeded on the top chamber of Transwell® Polycarbonate (PC) translucent 8-μm pore inserts (Corning, NY, USA) in 24-well plates at a density of 3x10^5^ cells per well in DMEM supplemented with 0.4% FBS. After 3 h, CXCL12 was added to the bottom chamber to the final concentration of 40 nM. The cells were incubated for an additional 20 h in a 37°C/5% CO_2_ incubator. This duration was chosen based on prior work by others (52) as reference, with further optimization in our own hands. On the following day, the supports were placed in a clean well containing 4% PFA for 1 h at room temperature, stained with crystal violet for 1 h, and washed three times in PBS. Cells on the upper side of the filters were removed with cotton-tipped swabs, and the number of migrated cells on the bottom side of the filter was counted in five randomly chosen fields at 200X magnification and averaged. All experiments were performed in triplicate, and each experiment was repeated at least three times.

### Stability Analysis

The mutation-induced changes in stability of Gαi were calculated as differences in the free energy of folding between the WT Gαi and mutants. The calculations were carried out in ICM (Molsoft LLC), using the *_*mutantStabilityAve tool and either GPCR bound [PDB: 6cmo (69), 6ot0 (64)], GDP bound [PDB: 1bof (101), 1gdd (102)), Gβγ bound (PDB: 1gg2 (103)] GTP bound [PDB: 1gia (104)] Gαi conformations. Predicted ddG values were plotted as a bar graph, with higher/more positive values indicating a higher degree of predicted destabilizing effect of the mutation on the given conformation.

### Confocal Immunofluorescence Microscopy

HeLa cells were plated on fibronectin coated coverslips at 500 K per well in a 6 well plate. Next day, cells were transfected with various Gαi3 mutants. 48 h post transfection, the coverslips were washed with PBS, treated with 4% paraformaldehyde for 30 min at RT, and again washed with PBS (3 times). The coverslips were blocked in blocking buffer (PBS, 0.1% Triton TX-100, and 0.5% BSA) in for 30 min and then incubated in a 1:100 dilution of the mouse monoclonal primary Gαi3 antibody (sc-365422, Santa Cruz, USA) in blocking buffer overnight in a moist chamber at 4^ο^C. The following day the coverslips were rinsed three times in PBS, each time allowing for a 15 min incubation. The coverslips were then treated with AlexaFluor488-conjugated secondary antibody (1:500 dilution) and DAPI (stock of 1 mg/ml diluted 1:2000, final concentration of 500 ng/mL) in blocking buffer for 45 min at room temperature, followed by rinsing them three times in PBS, each lasting 15 min. Coverslips were then blotted dry and mounted on a microscope slide with 5 µL Prolong Glass Antifade reagent (Invitrogen, USA). Transparent nail polish was applied for sealing the coverslips on the microscope slide and hold them in position. Visualization was done using SPE confocal microscope (Leica, Germany) using a 63X oil objective using 488nm and 405nm lasers for excitation. Around 10 images were collected for each construct. The settings were optimized, and the final images scanned with line-averaging of 3. All images were processed using ImageJ software (NIH) and assembled for presentation using Photoshop and Illustrator software programs (Adobe).

### Cytosol and Membrane Fractionation

HeLa cells were plated in 10 cm dishes at 2x10^6^ cells per dish in DMEM supplemented with 10% FBS. The next day cells were transfected with 8 μg/dish of total plasmid DNA’s of Gαi3(C351I) or phosphomutants using polyethylenimine (PEI), as described previously (105). Forty-eight hours after transfections, cells were rinsed once with PBS and lifted using a cell scraper in a homogenization buffer [25mM HEPES-KOH, 250 mM sucrose, 1 mM EDTA, pH 7.4] supplemented with protease and phosphatase inhibitors. Approximately 150 µl of homogenization buffer was used per 10 cm dish of cells. A 30-1/2 G syringe was used to break the cells by passing them through the needle ∼40 times (10 times/set with a 5 min rest time in-between each set, for a total of 4 sets). The suspension was then spun down at 500 g for 10 min (to discard the nuclear pellet), after which the supernatant was collected and transferred into a 1.7 ml thick-walled ultracentrifuge tube and ultracentrifuged at 100,000xg for 60 min. The resultant supernatant fraction was collected as the cytosolic fraction. The resultant pellet (crude membrane fraction) was rinsed further with homogenization buffer and centrifuged at 100,000xg for 30 min, and the pellet was subsequently resuspended in lysis buffer [20 mM HEPES, pH 7.2, 5 mM Mg-acetate, 125 mM K-acetate, 0.4% Triton X-100, 1 mM DTT, supplemented with sodium orthovanadate (500 µM), phosphatase (Sigma) and protease (Roche) inhibitor cocktails]. The proteins were quantified and 10% of each fraction, membrane and cytosol, were loaded on a 12% SDS PAGE gel and immunoblotted with Gαi3 antibody (11641-1-AP, ProteinTech, USA) and pan Gβγ antibody (sc 378, Santa Cruz, USA).

### Quantitative Immunoblotting

For immunoblotting, proteins were fractionated by SDS-PAGE and transferred to 0.4 µm PVDF membranes (Millipore, MA) prior to immunoblotting. Membranes were blocked with 5% nonfat milk dissolved in PBS (blocking buffer) before incubation with primary antibodies. Primary antibodies were prepared in blocking buffer containing 0.1% Tween-20 and incubated with blots overnight at 4^ο^C with rocking. After incubation, blots were washed and incubated with secondary antibodies for one hour at room temperature, washed again, and imaged a dual-color Li-Cor Odyssey imaging system (Li-Cor, USA). All Odyssey images were processed using Image J software (NIH) and assembled for presentation using Photoshop and Illustrator software programs (Adobe).

### Image Processing

All images were processed on ImageJ software (NIH) or FlowJo software and assembled into figure panels using Photoshop and Illustrator (Adobe Creative Cloud). All graphs were generated using GraphPad Prism v9.

### Statistical Analysis and Replicates

All experiments were repeated at least three times, and results were presented either as average ± S.E.M or with boundaries for 95% confidence interval. Statistical significance was assessed using F-tests for model fitting and regression analysis, and one-way analysis of variance (ANOVA) including a Tukey’s test for multiple comparisons. In general, we showed exact p values throughout the work. When depicted as “*”, they denote *p < 0.05, **p < 0.01, ***p < 0.001, ****p < 0.0001.

## Supplementary Materials

- Supplementary Materials and Methods

`o` Key Resource Table
- Supplementary Figures and legends (6)

`o` Figure S1: Functional validation of the Gαi3-FLAG construct.
`o` Figure S2: An alignment of the residues in Gαi that are phosphorylated with the corresponding residues in other Gα subunits across species.
`o` Figure S3: Effect of phosphomimicking mutations in Gαi3 on CXCL12-stimulated Gβγ activation downstream of CXCR4.
`o` Figure S4: Effect of phosphomimicking mutations in Gαi3 on basal and CXCL12-stimulated suppression of cAMP downstream of CXCR4.
`o` Figure S5: Effect of phosphomimicking and non-phosphorylatable mutants of Y320 on GiPCR recruitment to Gαi.
`o` Figure S6: Predicted impact of phosphomimicking or non-phosphorylatable mutations on the stability of Gαi in various conformations and complex compositions.
- Supplementary Tables and legends (6)

`o` Table S1: Statistical comparisons (ANOVA with Tukey’s multiple comparisons test) between the ability of indicated Gαi-mutants in **Fig 2e, f** to sequester Gβγ at baseline.
`o` Table S2: Statistical comparisons (ANOVA with Tukey’s multiple comparisons test) between the ability of indicated Gαi-mutants in **Fig 2g, h** to trigger trimer dissociation and release of Gβγ upon CXCL12 stimulation.
`o` Table S3: Statistical comparisons (ANOVA with Tukey’s multiple comparisons test) between the ability of indicated Gαi-mutants in **Fig 3c, d** to suppress cellular cAMP upon CXCL12 stimulation.
`o` Table S4: Model fitting and statistical hypothesis testing for mini-Gαs/i-WT, Y320E and Y320F studies shown in **Fig 4e, f**.
`o` Table S5: Statistical comparisons of Lineweaver-Burk (LB) plots for mini-Gαs/i-WT, Y320E and Y320F studies shown in **Supplementary Figure 5b-c**.
`o` Table S6: Statistical comparisons (ANOVA with Tukey’s multiple comparisons test) between the ability of indicated Gαi-mutants in **Fig 5c, d** to suppress chemotactic cell migration across a CXCL12 gradient.
- Supplementary Data Files

`o` Data File 1: Multiple Reaction Monitoring-based phosphorylation site analysis of immunopurified Gαi3-FLAG.
`o` Data File 2: Summary of phosphopeptides identified by targeted MRM.
`o` Data File 3: Source data for the HeatMap displayed in Figure 1e.

## ACKNOWLEDGMENTS

The authors thank A. Spiegel (National Institutes of Health, Bethesda, MD), Paul Insel and Tracy M. Handel (UC San Diego, CA, USA), Nevin A. Lambert (Augusta University, GA, USA), Michel Bouvier (Université de Montréal, Canada) and Stephan Lanier (University of South Carolina) for their generosity in sharing receptor and reporter constructs, and Tracy M. Handel (UC San Diego, CA, USA) for sharing access to the Guava benchtop flow cytometer.

## FUNDING

This paper was supported by the NIH (CA238042, AI141630, CA100768 and CA160911 to P.G., R21 AI149369, R01 GM136202, R21 AI156662, and R01 AI161880 to I.K.). S.S was supported through The American Association of Immunologists (AAI) Intersect Fellowship Program for Computational Scientist and Immunobiologists.

## AUTHOR CONTRIBUTIONS

I.K and P.G conceptualized the project; S.R, with assistance from A.S, performed experiments; I.K carried out all the computational analyses; M.G and P.G carried out the *in cellulo* phosphorylation and LC/MS studies. S.R, S.S, I.K and P.G prepared figures for data visualization. SS provided expertise in statistical analysis. S.R, I.K and P.G wrote the original draft of the manuscript. All authors provided input and edited and revised the manuscript. All co-authors approved the final version of the manuscript. I.K and P.G coordinated and supervised all parts of the project and administered the project.

## COMPETING INTERESTS

The authors declare no competing interests.

## DATA AND MATERIALS AVAILABILITY

All data for this research are included in the manuscript and in Supplementary Materials.

## Notes

### Competing Interest Statement

The authors have declared no competing interest.

### Summary of Updates

Revised throughout text, figures, legends. Addition of new author (Sinha).

