## Supplementary Online Materials for "Phosphorylation of Gαi shapes canonical Gα(i)βγ/GPCR signaling"

- Supplementary Materials and Methods
    - Key Resource Table
- Supplementary Figures and legends (6)
    - Figure S1
    - Figure S2
    - Figure S3
    - Figure S4
    - Figure S5
    - Figure S6

- Supplementary Tables and legends (6)
    - Table S1
    - Table S2
    - Table S3
    - Table S4
    - Table S5
    - Table S6

- Supplementary Data Files
    - Data File 1
    - Data File 2
    - Data File 3

### Supplementary Material and Methods

#### KEY RESOURCES TABLE

| REAGENT or RESOURCE | SOURCE | IDENTIFIER |
|---|---|---|
| **BIOLOGICAL SAMPLES AND CELL LINES** | | |
| **CELL LINE** | Source | Cat# |
| **HeLa parental** | ATCC | ATCC® CCL-2 |
| **Cos 7** | ATCC | ATCC® CRL-1651 |
| **PLASMIDS AND CONSTRUCTS** | | |
| **HA-CXCR4-Rluc3** | *Tracy Handel* | (87) |
| **CAMYEL (modified sensor)** | *Paul Insel and Tracy Handel* | (68, 88) |
| **masGRK3ct-RLuc8** | *Nevin Lambert* | (89) |
| **mVenus-hGBB1** | *Nevin Lambert* | (89) |
| **mVenus-hGBG2** | *Nevin Lambert* | (89) |
| **NES-mVenus-miniGαs/i** | *Nevin Lambert* | (87) |
| **Rat Gαi3 (untagged)** | *A. Spiegel* | (90) |
| **Gαi3(C351I)** | *This work* | |
| **Gαi3-S44D(C351I)** | *This work* | |
| **Gαi3-S151D(C351I)** | *This work* | |
| **Gαi3-S47D-T48E(C351I)** | *This work* | |
| **Gαi3-YY154-155EE(C351I)** | *This work* | |
| **Gαi3-Y320E(C351I)** | *This work* | |
| **Gαi3-Y320F(C351I)** | *This work* | |
| **Gαi3-YYY154-155-320-EEE(C351I)** | *This work* | |
| **Gαi3-YYY154-155-320-EEF(C351I)** | *This work* | |
| **Gαi3-S44D-S47D-T48E-Y154E-Y155E(C351I)** | *This work* | |
| **GST-tagged GAIP** | | (91, 92) |
| **GST-tagged GoLoco region of AGS3** | *Stephan Lanier* | (35) |
| **His-tagged GIV-C terminus short** | | (92) |
| **CHEMICALS, KITS, RECOMBINANT PROTEINS** | | |
| **FETAL BOVINE SERUM, NEW ZEALAND ORIGIN** | Sigma | F8067 |
| **DMEM** | Cytiva | SH30243.FS |
| **OptiMEM** | Thermo Scientific | 11058021 |
| **Antibiotic-Antimycotic** | Gibco | 15240062 |
| **Accutase** | BioLegend | 423201 |
| **Trypsin-EDTA** | Genesee Scientific | 25-510 |

| | | |
|---|---|---|
| **96 well plate, white/clear bottom** | Falcon | 353296 |
| **TransIT-X2 transfection reagent** | Mirus | MIR6000 |
| **Pertussis Toxin** | VWR | 102946-454 |
| **Coelenterazine-H** | RPI corporation | C615000.001 |
| **Forskolin** | Sigma | F6886 |
| **IBMX** | Sigma | I5879 |
| **Protease inhibitor cocktail** | Roche | 11 873 580 001 |
| **Tyr phosphatase inhibitor cocktail** | Sigma-Aldrich | P5726 |
| **Ser/Thr phosphatase inhibitor cocktail** | Sigma-Aldrich | P0044 |
| **PVDF Transfer Membrane** | Thermo Scientific | 88518 |
| **Guava Cell Cycle Reagent (ICF)** | Millipore Sigma | 4700-0160 |
| **8 µm pore Polycarbonate Transwell plate** | Corning | 3422 |
| **Paraformaldehyde 16%** | Electron Microscopy Biosciences | 15710 |
| **Prolong Glass** | Thermo Fisher Scientific | P36980 |
| **PEI** | Polysciences INC Research Chem | 239661 |
| **Gibco fibronectin human protein, native** | Thermo Scientific | PHE0023 |
| **ANTIBODIES** | | |
| **Mouse monoclonal anti-α-tubulin (B-7)** | Santa Cruz Biotechnology | sc-5286 |
| **Mouse monoclonal anti-FLAG** | Millipore Sigma | MAB3118 |
| **Rabbit polyclonal pan anti-Gβ** | Santa Cruz Biotechnology | sc-378 |
| **Mouse monoclonal anti-Gαi3** | Santa Cruz Biotechnology | SC365422 |
| **Rabbit polyclonal anti-Gαi3** | ProteinTech | 11641-1-AP |
| **Mouse monoclonal anti-β1-subunit of Na+/K+-transporting ATPase (ATP1B1)** | Santa Cruz Biotechnology | sc-21713 |
| **Mouse monoclonal anti-GAPDH** | Santa Cruz Biotechnology | sc-47724 |
| **Anti-HA antibody, APC-conjugated** | *Miltenyi Biotech* | 130-123-639 |
| **Goat anti-Mouse IgG, Alexa Fluor 488 conjugated** | ThermoFisher Scientific | A11017 |
| **IRDye 800CW Goat anti-Mouse IgG Secondary** | LI-COR Biosciences | 926-32210 |
| **IRDye 680RD Goat anti-Rabbit IgG Secondary** | LI-COR Biosciences | 926-68071 |
| **SOFTWARE** | | |

| | | |
|---|---|---|
| **ImageJ** | National Institute of Health | https://imagej.net/Welcome |
| **FlowJo** | FlowJo, LLC | https://www.flowjo.com |
| **Prism** | GraphPad | https://www.graphpad.com/scientific-software/prism/ |
| **LAS-X** | Leica | www.leica-microsystems.com/products/microscope-software/p/leica-las-x-ls |
| **Molsoft ICM Pro** | Molsoft, LLC | https://www.molsoft.com/index.html |
| **Illustrator** | Adobe | https://www.adobe.com/products/illustrator.html |
| **ImageStudio Lite** | LI-COR | https://www.licor.com/bio/image-studio-lite/ |
| **Biorender** | Biorender | https://www.biorender.com/ |
| **BioEdit** | BioEdit 7.7 | https://bioedit.software.informer.com/ |

### Supplementary figures and legends

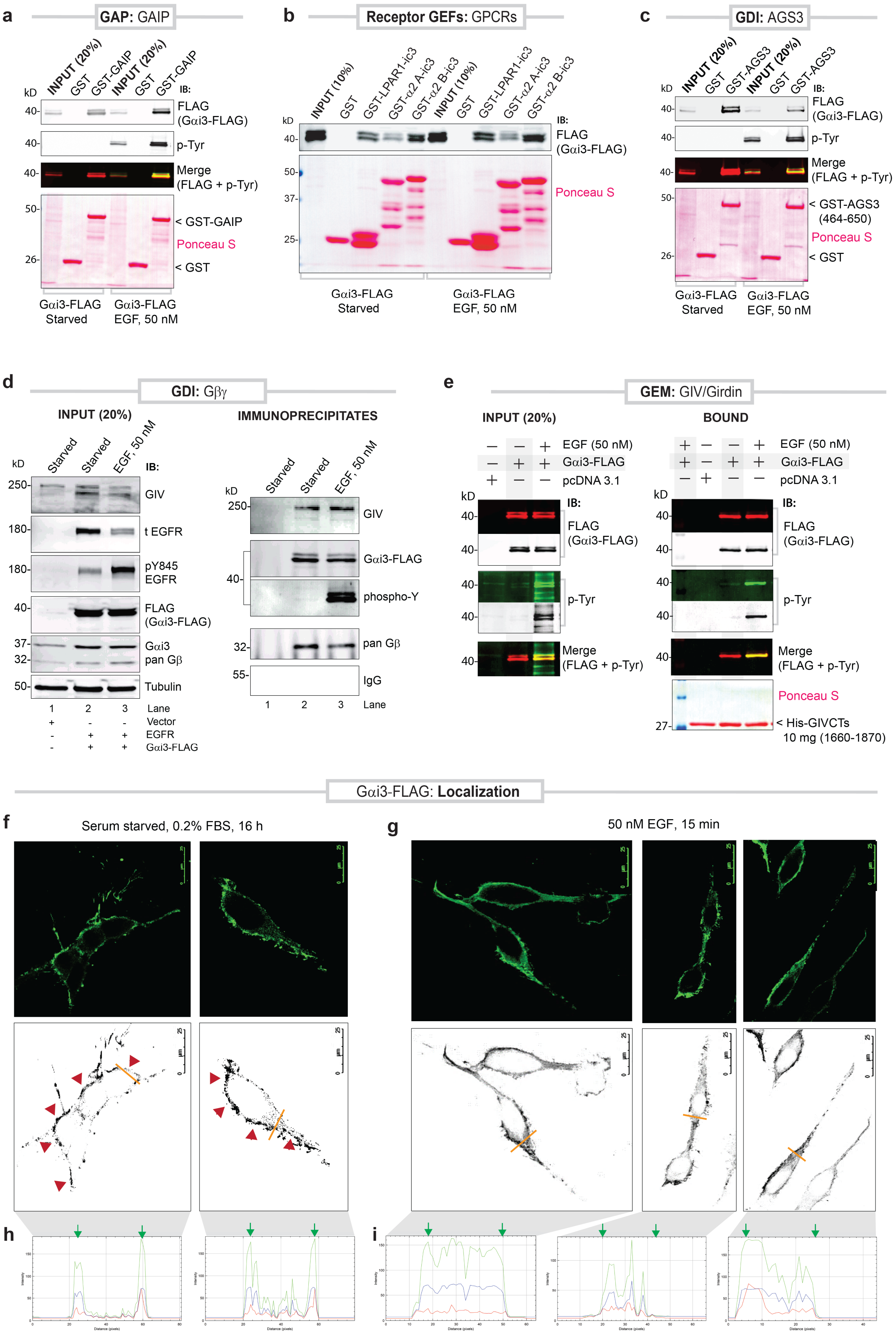


**Figure S1**. **Functional validation of the Gαi3-FLAG construct.**

a. Pull down assays with GST-tagged GAIP (~5 µg) or GST alone (negative control) and lysates of serum-starved or EGF (50 nM) stimulated Cos7 cells expressing Gαi3-FLAG were analyzed by for bound Gαi3 by dual-color immunoblotting (IB). Anti-FLAG mAb (total Gαi3; red) and anti-p-Tyr mAb (tyrosine phosphorylated Gαi3; green) channels are displayed as individual grayscale images and as colored 'merged' panels. Ponceau stained membrane confirms equal loading of GST proteins.

b. Pull down assays with GST-tagged 3$^{rd}$ intracellular loop (icl3; ~15 µg) of the indicated GPCRs or GST alone (negative control) and lysates of serum-starved or EGF (50 nM) stimulated Cos7 cells expressing Gαi3-FLAG were analyzed by for bound Gαi3. Ponceau stained membrane shows the various GST-tagged proteins.

c. Pull down assays with GST-tagged GoLoco region of AGS3 (~5 µg) or GST alone (negative control) and lysates of serum-starved or EGF (50 nM) stimulated Cos7 cells expressing Gαi3-FLAG were analyzed by for bound Gαi3 by dual-color immunoblotting (IB). Anti-FLAG mAb (total Gαi3; red) and anti-p-Tyr mAb (tyrosine phosphorylated Gαi3; green) channels are displayed as individual grayscale images and as colored 'merged' panels. Ponceau stained membrane confirms equal loading of GST proteins.

d. Immunoprecipitation assays were carried out using a monoclonal antibody against the FLAG epitope and equal aliquots of lysates of Cos7 cells exogenously expressing Gαi3-FLAG and starved and EGF-stimulated exactly as in **Figure 1a**. Bound proteins were analyzed for Gαi3 and Gβγ by immunoblotting (IB). EGFR activation in cell lysates (left) and Gαi3 phosphorylation in immunoprecipitated complexes (right) were confirmed by immunoblotting for pYEGFR and pan-pTyr using monoclonal antibodies.

e. Pull down assays with His-tagged GIV-C terminus short (CTs) (~10 µg) and lysates of serum-starved or EGF (50 nM) stimulated Cos7 cells expressing Gαi3-FLAG were analyzed by for bound Gαi3 by dual-color immunoblotting (IB). Anti-FLAG mAb (total Gαi3; red) and anti-p-Tyr mAb (tyrosine phosphorylated Gαi3; green) channels are displayed as individual panels in color and as grayscale images and as 'merged' panels. Ponceau stained membrane confirms equal loading of His-GIVCT proteins.

f-i. Serum-starved or EGF-stimulated HeLa cells transiently transfected with Gαi3-FLAG were fixed, stained and analyzed by confocal immunofluorescence microscopy. Representative images are displayed. *Upper panels*: green pixels show Gαi3 visualized using anti-FLAG mAb. *Lower panels*: grayscale versions of upper panels (red arrowheads indicate peripheral membrane localization. Scale bars: 25 µm. Orange lines in panel g indicate the regions of interest analyzed for signal intensity using ImageJ (RGB plot graphics; panels h-i). Green arrows indicate the presumed plasma membrane at the cell border. Unlike the two distinct intensity peaks in panel h, we observe a more diffused signal in panels j.

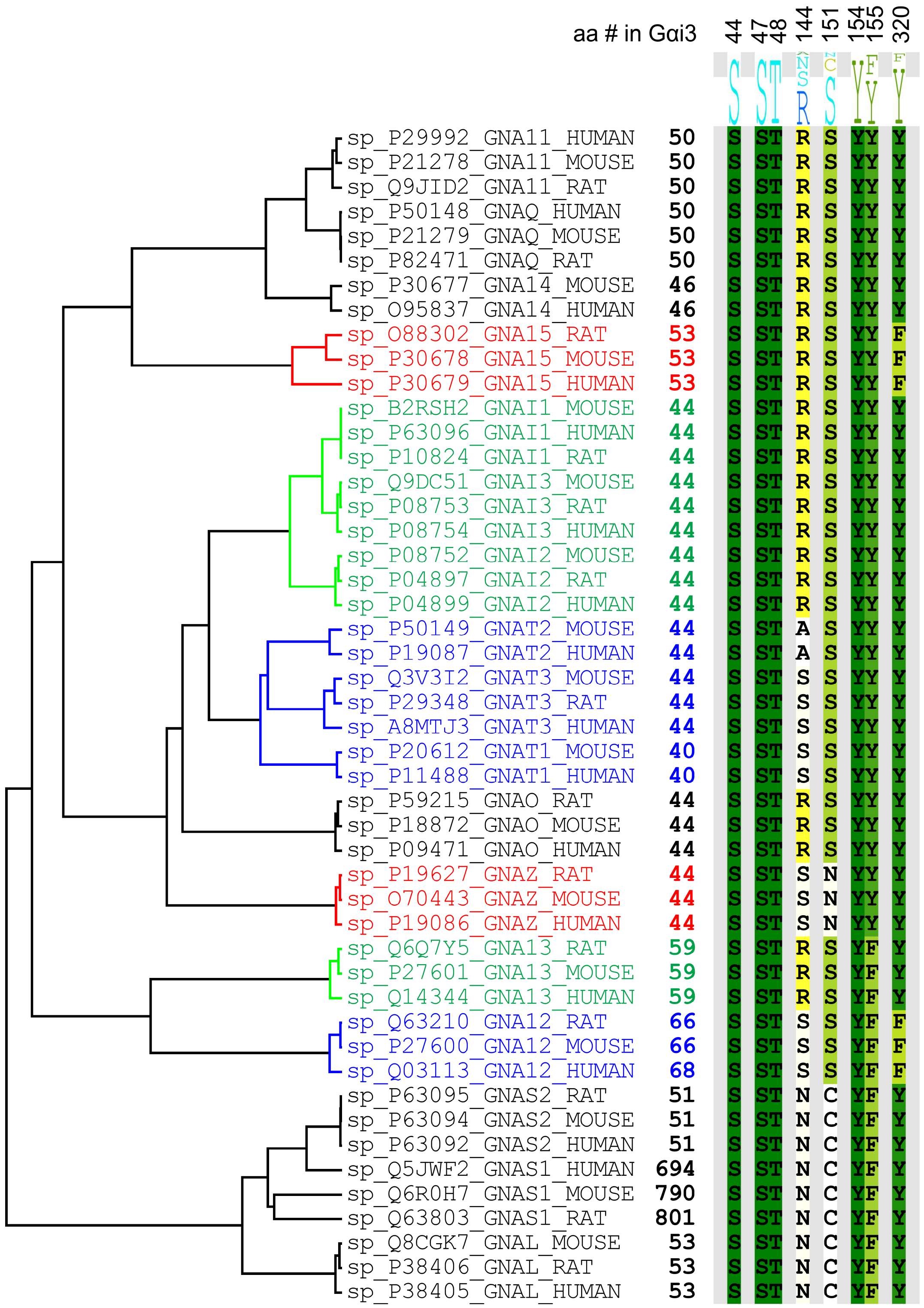


**Figure S2**. **An analysis of the degree of conservation of the phosphosites identified in this work**. An alignment of human, mouse, and rat Gα proteins of different subtypes, projected to the amino-acid positions studied in this work. The alignment illustrates the absolute conservation of $Ser^{44}$, $Ser^{47}$, $Thr^{48}$ and $Y^{154}$ across all subtypes, as well as the possible aa substitutions for $S^{144}$ (Arg, Ala, and Asn), $S^{151}$ (Asn and Cys), $Y^{155}$ (Phe) and $Y^{320}$ (Phe).

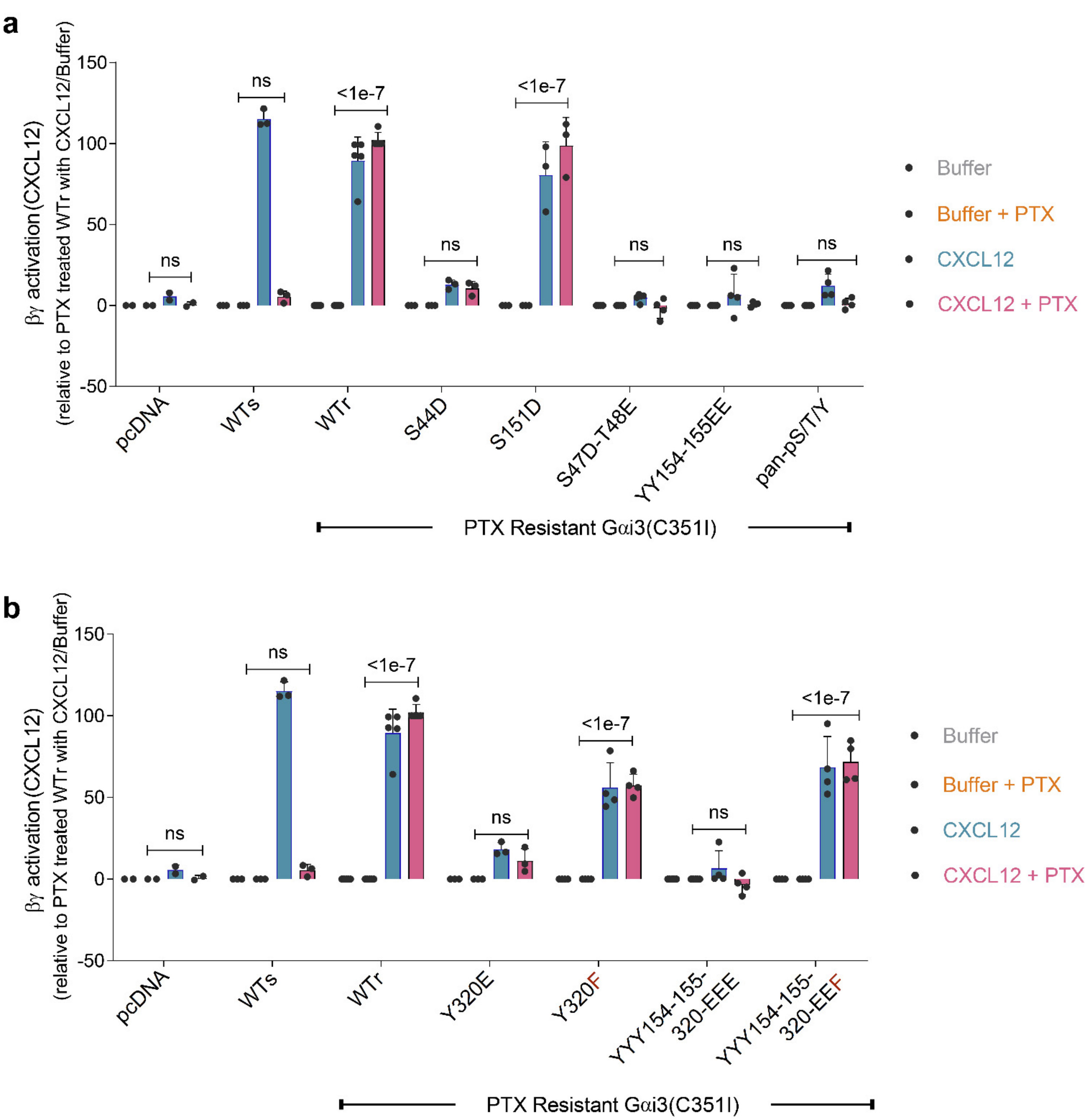


**Figure S3**. **Effect of phosphomimicking mutations in Gαi3 on CXCL12-stimulated Gβγ activation downstream of CXCR4.**

a-b. Bar graphs display the % Gβγ activation upon ligand stimulation observed in the case of each indicated mutant, as assessed in 4 conditions i.e., with and without PTX pre-treatment and with and without CXCL12 stimulation. Results are expressed as normalized values compared to the observed activity in PTX-resistant Gαi3 (WTr, set at 100%) and to PTX-sensitive Gαi3 (WTs, set at 0%) in the PTX-treated condition. *P* values were determined by two-way ANOVA with Tukey's multiple comparison's test. Statistics are displayed comparing the buffer/PTX and CXCL12/PTX conditions. Error bars represent ± S.E.M; n=4 independent experiments, each with 3 technical replicates.

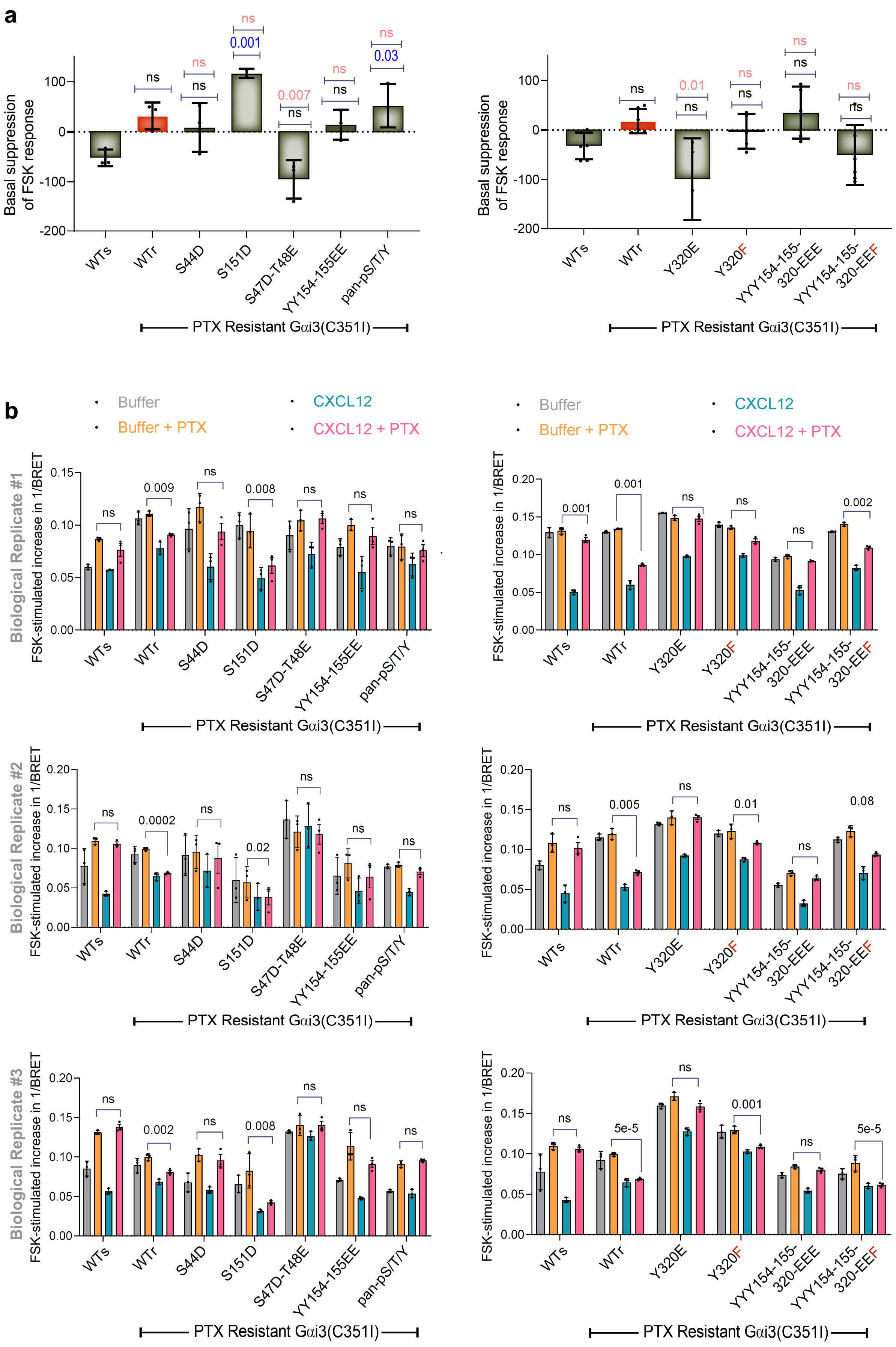


**Figure S4. Effect of phosphomimicking mutations in Gαi3 on basal and CXCL12-induced suppression of FSK-stimulated cAMP downstream of CXCR4.**

a. Results of the BRET-based CAMYEL (*cAM*P sensor using *Y*FP-*E*pac-R*L*uc) assay for assessing cellular cAMP are shown. Bar graphs display the % suppression of forskolin- (FSK-) cAMP observed in the case of each mutant at baseline prior to CXCL12 stimulation. Results are displayed as fold change compared to the observed basal activity in the PTX-sensitive Gαi3 (WTs, set at 0%) and PTX-resistant Gαi3 (WTr, set at 100%) in the PTX-treated condition.

b. Bar graphs display the FSK-induced change in the inverse BRET in HeLa cells transiently expressing the of the CAMYEL cAMP sensor and the indicated Gαi mutants, pretreated or not with PTX, and stimulated or not with CXCL12. A decrease in BRET in the presence of the ligand (CXCL12 + PTX) compared to control (Buffer + PTX) implies suppression of FSK-stimulated cAMP. The top, middle, and lower panels represent three distinct biological repeats. *P* values were determined by two-way ANOVA with Tukey's multiple comparison's test. Statistics are displayed comparing the buffer/PTX and CXCL12/PTX conditions. Error bars represent ± S.E.M; n=4 independent experiments, each with 3 technical replicates.

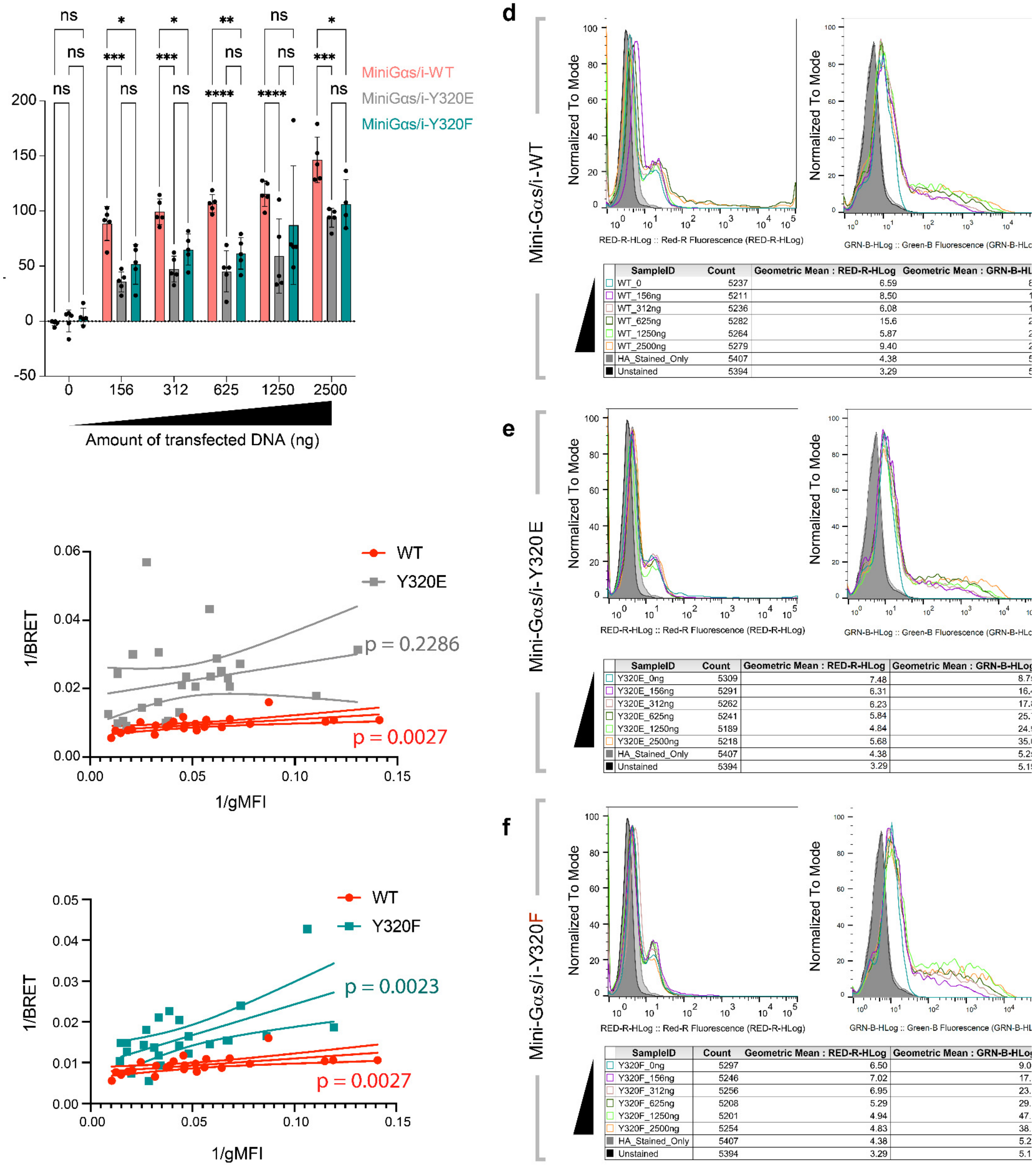


**Figure S5. Effect of phosphomimicking and non-phosphorylatable mutants of Y320 on GiPCR recruitment to Gαi.**

a. Bar graphs showing the CXCL12 response (BRET ratio normalized to mini-Gαs/i-WT), for the increasing amount of the acceptors (mVenus-mini-Gαs/i-WT, mVenus-mini-Gαs/i-320E and mVenus-mini-Gαs/i-320F). The use of *, **, ***, **** mark statistical significance; ns, not significant; $p<.05$; $p<.01$; $p<.001$; $p<.0001$.

b, c. Lineweaver-Burk (LB) plots of 1/BRET (y axis) vs 1/gMFI (x axis) with 95% confidence boundaries (interrupted lines) for the comparisons between mini-Gαs/i-WT vs mini-Gαs/i-Y320E (b) and mini-Gαs/i-WT vs mini-Gαs/i-Y320F (c).

d- f. Titration of the acceptor Venus-tagged Mini Gαs/i expression, as confirmed by flow cytometry. Distributions of mVenus (GRN-B-HLog, indicative of the expression of mVenus-mini-Gαs/i, the BRET acceptor) and anti-HA-APC (RED-R-HLog, reflective of surface expression of HA-CXCR4-RLucII, the donor) fluorescence intensities in samples used for the acceptor titration BRET experiments in **Figure 4**. Un-transfected HeLa cells that were unstained or stained with anti-HA-APC were used as negative controls.

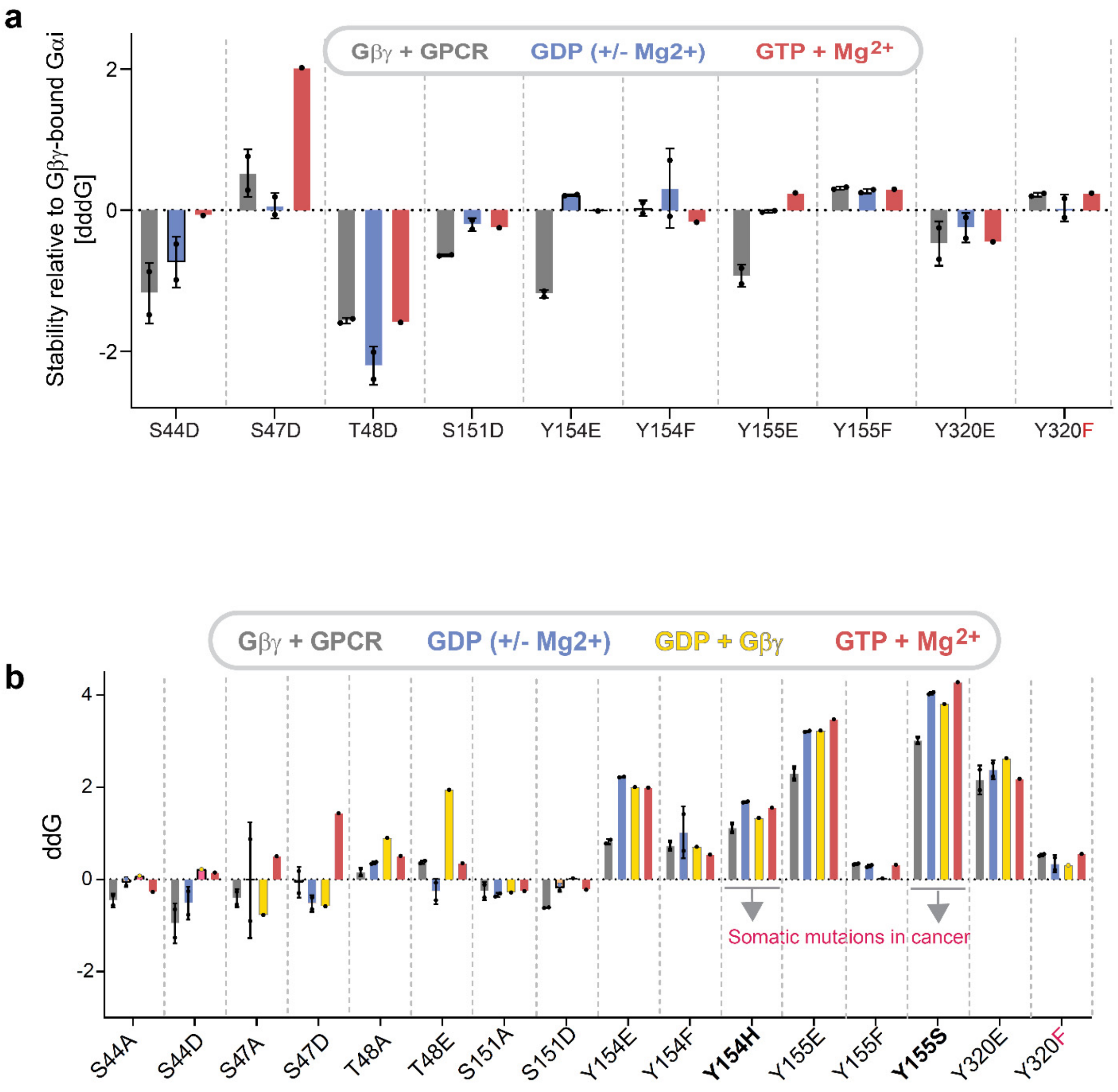


**Figure S6. Predicted impact of phosphomimicking or non-phosphorylatable mutations on the stability of Gαi in various conformations and complex compositions.**

a. Effects of various mutations on the stability of Gαi3 in Gβγ+GPCR-, GDP-, and GTP-bound states relative to the Gβγ+GDP-bound state. Higher / positive bars indicate that the given complex is destabilized to a larger degree and therefore is less favorable than the Gβγ+GDP-bound state, negative bars indicate the opposite.

b. Bar graphs displays the computationally predicted effects of various mutations on the stability of Gαi3 in various conformations and complex compositions, including a complex with Gβγ+GPCR (PDB:6cmo,6ot0), GDP (+/-$Mg^{2+}$; PDB:1bof, 1gdd), GDP+Gβγ (PDB:1gg2), and GTP+$Mg^{2+}$ (PDB:1gia). Bar height indicates delta delta G [ddG], i.e., the predicted change in the change in Gibbs free energy of folding; higher / more positive bars reflect a larger degree of destabilization by the respective mutation while negative numbers indicate the opposite. The analysis also includes somatic mutations in cancers (COSMIC; https://cancer.sanger.ac.uk/cosmic) that affect those sites, Y154H and Y155S, which were characterized in a prior work (5).

### Supplementary Tables

**Table S1**. Statistical comparisons (ANOVA with Tukey's multiple comparisons test) between the ability of indicated Gαi-mutants in **Fig 2e, f** to sequester Gβγ at baseline.

| Tukey's multiple comparisons test | Mean Diff. | 95.00% CI of diff. | Significant [Yes/No] | Summary | Adj. p Value |
|---|---|---|---|---|---|
| **Y320E vs. Y320F** | -46.34243 | -81.04 to -11.63 | Yes | **** | 1.40e-03 |
| **Y320E vs. YYY154-155-320-EEE** | 44.06463 | 9.35 to 78.77 | Yes | **** | 3.00e-03 |
| **Y320E vs. YYY154-155-320-EEF** | -47.41751 | -84.52 to -10.31 | Yes | **** | 2.8e-3 |
| **Y320E vs. YY154-155EE** | 42.24582948 | 7.53 to 76.95 | Yes | **** | 5.4e-03 |
| **Y320F vs. YYY154-155-320-EEE** | 90.40706 | 58.27 to 122.54 | Yes | **** | <1 e-7 |
| **Y320F vs. YYY154-155-320-EEF** | -1.075076 | -35.78 to 33.63 | No | ns | 1 |
| **Y320F vs. YY154155EE** | 88.58826069 | 56.45 to 120.72 | Yes | **** | <1 e-7 |
| **YYY154-155-320-EEE vs. YYY154-155-320-EEF** | -91.48214 | -126.18 to -56.77 | Yes | **** | <1 e-7 |
| **YYY154-155-320-EEE vs. YY154155EE** | -1.818801475 | -33.95 to 30.31 | No | ns | 1 |
| **YYY154-155-320-EEF vs. YY154155EE** | 89.66333668 | 54.95 to 124.37 | Yes | **** | <1 e-7 |

**Table S2**. Statistical comparisons (ANOVA with Tukey's multiple comparisons test) between the ability of indicated Gαi-mutants in **Fig 2g, h** to trigger trimer dissociation and release of Gβγ upon CXCL12 stimulation.

| Tukey's multiple comparisons test | Mean Diff. | 95.00% CI of diff. | Significant [Yes/No] | Summary | Adj. p Value |
|---|---|---|---|---|---|
| **Y320E vs. Y320F** | -46.17309373 | -67.03 to -25.31 | Yes | **** | 6.98e-07 |
| **Y320E vs. YYY154-155-320EEE** | 14.73951556 | -6.11 to 35.59 | No | ns | 0.37516 |
| **Y320E vs. YYY154-155-320-EEF** | -56.21693965 | -78.51 to -33.91 | Yes | **** | <1 e-7 |
| **Y320E vs. YY154-155EE** | 10.52163 | -10.33 to 31.37 | No | ns | 0.4093172 |
| **Y320F vs. YYY154-155-320EEE** | 60.91261 | 41.60 to 80.22 | Yes | **** | <1 e-7 |
| **Y320F vs. YYY154-155-320-EEF** | -10.04384592 | -30.90 to 10.81 | No | ns | 0.85072385 |
| **Y320F vs. YY154-155EE** | 56.69472 | 37.38 to 76.00 | Yes | **** | <1 e-7 |
| **YYY154-155-320EEE vs. YYY154-155-320EEF** | -70.9564552 | -91.81 to -50.09 | Yes | **** | <1 e-7 |
| **YYY154-155-320EEE vs. YY154-155EE** | -4.217889 | -23.52 to 15.09 | No | ns | 0.99964353 |
| **YYY154-155-320EEF vs. YY154-155EE** | 66.73856631 | 45.88 to 87.59 | Yes | **** | <1 e-7 |

**Table S3**. Statistical comparisons (ANOVA with Tukey's multiple comparisons test) between the ability of indicated Gαi-mutants in **Fig 3c, d** to suppress cellular cAMP upon CXCL12 stimulation.

| Tukey's multiple comparisons test | Mean Diff. | 95.00% CI of diff. | Significant [Yes/No] | Summary | Adj. p Value |
|---|---|---|---|---|---|
| **Y320E vs. Y320F** | -54.1149 | -131.30 to 23.07 | No | ns | 0.251567 |
| **Y320E vs. YYY154-155-320EEE** | -6.0113 | -87.37 to 75.35 | No | ns | 0.999372 |
| **Y320E vs. YYY154-155-320-EEF** | -94.6507 | -171.83 to -17.46 | Yes | * | 0.012515 |
| **Y320E vs. YY154-155EE** | -77.1041 | -158.46 to 4.25 | No | ns | 0.067927 |
| **Y320F vs. YYY154-155-320EEE** | 48.10362 | -29.08 to 125.29 | No | ns | 0.356119 |
| **Y320F vs. YYY154-155-320-EEF** | -40.5358 | -113.30 to 32.23 | No | ns | 0.462631 |
| **Y320F vs. YY154-155EE** | -22.9892 | -100.17 to 54.19 | No | ns | 0.890625 |
| **YYY154-155-320EEE vs. YYY154-155-320EEF** | -88.6394 | -165.82 to -11.45 | Yes | * | 0.020325 |
| **YYY154-155-320EEE vs. YY154-155EE** | -71.0928 | -152.45 to 10.272 | No | ns | 0.103238 |
| **YYY154-155-320EEF vs. YY154-155EE** | 17.5466 | -59.64 to 94.73 | No | ns | 0.955591 |

**Table S4:** Model fitting and statistical hypothesis testing for mini-Gαs/i-WT, Y320E and Y320F studies shown in **Fig 4e, f**.

| **Mini-Gαs/i-WT vs Mini-Gαs/i-Y320E** | | | | |
|---|---|---|---|---|
| **Best fit** | **Best-fit values** | **Mini-Gαs/i-WT** | **Mini-Gαs/i-Y320E** | **Global** |
| | **Bmax** | 148.6 | 93.69 | 112.8 |
| | **Kd** | 7.102 | 16.69 | 7.785 |
| **Global fit comparison** | **R squared** | 0.5752 | -0.1004 | 0.5164 |
| | **Sum of Squares** | 28496 | 33975 | 62470 |
| **Null hypothesis** | One curve for all data sets | | | |
| **Alternative hypothesis** | Different curve for at least one data set | | | |
| **P value** | <0.0001 | | | |
| **Conclusion (alpha = 0.05)** | **Reject null hypothesis** | | | |
| **Preferred model** | Different curve for at least one data set | | | |
| **F (DFn, DFd)** | 54.65 (2, 56) | | | |
| **Mini-Gαs/i-WT vs Mini-Gαs/i-Y320F** | | | | |
| **Best fit** | **Best-fit values** | **Mini-Gαs/i-WT** | **Mini-Gαs/i-Y320F** | **Global** |
| | **Bmax** | 148.6 | 112.9 | 125.7 |
| | **Kd** | 7.102 | 13.44 | 8.109 |
| **Global fit comparison** | **R squared** | 0.7338 | 0.3253 | 0.6201 |
| | **Sum of Squares** | 17855 | 30152 | 48007 |
| **Null hypothesis** | One curve for all data sets | | | |
| **Alternative hypothesis** | Different curve for at least one data set | | | |
| **P value** | <0.0001 | | | |
| **Conclusion (alpha = 0.05)** | **Reject null hypothesis** | | | |
| **Preferred model** | Different curve for at least one data set | | | |
| **F (DFn, DFd)** | 20.69 (2, 55) | | | |

**Table S5:** Statistical comparisons of Lineweaver-Burk (LB) plots for mini-Gαs/i-WT, Y320E and Y320F studies shown in **Supplementary Figure 5b-c**.

| | | Mini-Gαs/i-WT | Mini-Gαs/i-Y320E | Mini-Gαs/i-Y320F |
|---|---|---|---|---|
| **Best-fit values** | **Slope** | 0.03449 | 0.09435 | 0.1509 |
| | **Y-intercept** | 0.007563 | 0.01776 | 0.009196 |
| | **X-intercept** | -0.2193 | -0.1883 | -0.06093 |
| | **1/slope** | 29 | 10.6 | 6.626 |
| **Goodness of Fit** | **R squared** | 0.3292 | 0.06237 | 0.3507 |
| | **Sy.x** | 0.001709 | 0.01105 | 0.005958 |
| **Significantly non-zero slope** | **F** | 11.29 | 1.53 | 11.88 |
| | **DFn, DFd** | 1, 23 | 1, 23 | 1, 22 |
| | **P value** | 0.0027 | 0.2286 | 0.0023 |
| | **Deviation from zero?** | Significant | Not Significant | Significant |

**Table S6**. Statistical comparisons (ANOVA with Tukey's multiple comparisons test) between the ability of indicated Gαi-mutants in **Fig 5c, d** to suppress chemotactic cell migration across a CXCL12 gradient.

| Tukey's multiple comparisons test | Mean Diff. | 95.00% CI of diff. | Significant [Yes/No] | Summary | Adj. p Value |
|---|---|---|---|---|---|
| **Y320E vs. Y320F** | -36.702 | -70.63920 to -2.764731 | Yes | * | 0.028955 |
| **Y320E vs. YYY154-155-320EEE** | -9.08575 | -40.50553 to 22.33402 | No | ns | 0.958853 |
| **Y320E vs. YYY154-155-320EEF** | -55.6047 | -87.02443 to -24.18488 | Yes | *** | 0.000229 |
| **Y320E vs. YY154155EE** | -12.8552 | -44.27496 to 18.56459 | No | ns | 0.823214 |
| **Y320F vs. YYY154-155-320EEE** | 27.61621 | -6.321023 to 61.55345 | No | ns | 0.159246 |
| **Y320F vs. YYY154-155-320EEF** | -18.9027 | -52.83992 to 15.03455 | No | ns | 0.545749 |
| **Y320F vs. YY154-155EE** | 23.84679 | -10.09045 to 57.78402 | No | ns | 0.290006 |
| **YYY154-155-320EEE vs. YYY154-155-320EEF** | -46.5189 | -77.93867 to -15.09912 | Yes | ** | 0.001743 |
| **YYY154-155-320EEE vs. YY154155EE** | -3.76943 | -35.18920 to 27.65035 | No | ns | 0.999625 |
| **YYY154-155-320EEF vs. YY154155EE** | 42.74947 | 11.32969 to 74.16925 | Yes | ** | 0.004093 |

### Supplementary Data Files (Uploaded as excel sheets)

**Data File 1**: Multiple Reaction Monitoring-based phosphorylation site analysis of immunopurified Gαi3-FLAG.

**Data File 2**: Summary of phosphopeptides identified by targeted MRM.

**Data File 3**: Source data for the HeatMap displayed in **Figure 1e.**
